# Diversification in evolutionary arenas – assessment and synthesis

**DOI:** 10.1101/636803

**Authors:** Nicolai M. Nürk, H. Peter Linder, Renske E. Onstein, Matthew J. Larcombe, Colin E. Hughes, Laura Piñeiro Fernández, Philipp M. Schlüter, Luis Valente, Carl Beierkuhnlein, Vanessa Cutts, Michael J. Donoghue, Erika J. Edwards, Richard Field, Suzette G.A. Flantua, Steven I. Higgins, Anke Jentsch, Sigrid Liede-Schumann, Michael D. Pirie

## Abstract

Understanding how and why rates of evolutionary diversification vary is a key issue in evolutionary biology, ecology, and biogeography, and the metaphorical concepts of adaptive radiation and evolutionary stasis describe two opposing aspects causing variation in diversification rates. Here we review the central concepts in the evolutionary diversification literature and synthesize these into a simple, general framework for studying rates of diversification and quantifying their underlying dynamics, which can be applied across clades and regions and across spatial and temporal scales. Our framework describes the diversification rate (*d*) as a function of the abiotic environment (*a*), the biotic environment (*b*) and clade-specific phenotypes or traits (*c*); thus *d*∼*a,b,c*. We refer to the four components (*a*–*d*) and their interactions collectively as the ‘Evolutionary Arena’. We outline analytical approaches to this framework and present a case study on conifers, for which we parameterise the general model. We also discuss three conceptual examples: the *Lupinus* radiation in the Andes in the context of emerging ecological opportunity and fluctuating connectivity due to climatic oscillations; oceanic island radiations in the context of island formation and erosion; and biotically driven radiations of the Mediterranean orchid genus *Ophrys*. The results of the conifer case study are consistent with the long-standing scenario that low competition and high rates of niche evolution promote diversification. The conceptual examples illustrate how using the synthetic Evolutionary Arena framework helps to identify and structure future directions for research on evolutionary radiations. In this way, the Evolutionary Arena framework promotes a more general understanding of variation in evolutionary rates by making quantitative results comparable between case studies, thereby allowing new syntheses of evolutionary and ecological processes to emerge.

---

> *“In reviewing the literature, we are struck that there is no one formula for developing a convincing hypothesis about diversification and its causes*.*”*
>
> — (Donoghue & Sanderson, 2015, p. 263)

## Introduction

Biologists have long been fascinated by the circumstances under which species diversification and trait disparification rates – evolutionary radiations – are accelerated. Studies in recent decades on evolutionary **radiations** (words in bold are in the Glossary) have produced a proliferation of terminology and new statistical approaches. These developments in (macro-) evolution are largely based on the adaptive radiation paradigm (Osborn, 1902; Simpson, 1944, 1953; Schluter, 2000), a metaphorical concept describing the evolution of a multitude of ecological forms from a single common ancestor. The paradigm, however, complicates quantitative comparisons of the trajectories and correlates of diversification between evolutionary lineages (species, clades) and among geographical regions, and does not address the circumstances under which evolutionary stasis or decline may occur. Here, we build on current theoretical foundations and propose a conceptual framework for the integrative study of shifts and stasis in diversification rates. It is not our aim to thoroughly review the literature on evolutionary radiations; rather, we provide an overview of recent developments, and integrate these into a framework that can in principle be quantified in all systems, from cellular to global spatial scales and spanning ecological to evolutionary time frames.

### A short history of diversification theory

Darwin, in sharp contrast to early-nineteenth-century dogma, envisioned evolution to be gradual, with small changes accumulating from generation to generation, eventually leading to species divergence (Orr, 2005). This gradualist view was soon challenged and seemingly contradicted by the fossil record, leading to the appreciation that rates of divergent evolution are uneven through time and among clades, sometimes generating species and ecomorphological diversity in evolutionary radiations (Mayr, 1954; Stanley, 1979), while at other times demonstrating long-term stasis (Eldredge & Gould, 1972; Gould & Eldredge, 1977; Flegr, 2010) or decline in diversity (Benton, 1995; Rohde & Muller, 2005).

The development of phylogenetic theory (Hennig, 1950, 1965) followed by the generation of massive DNA sequence datasets, increased computing power, and the proliferation of analytical methods (i.a., maximum likelihood, Felsenstein, 1973, 1981; Bayesian inference, Huelsenbeck, Ronquist, Nielsen, & Bollback, 2001; Bayesian molecular dating, Drummond, Ho, Phillips, & Rambaut, 2006; multispecies coalescence, Degnan & Rosenberg, 2006; Edwards, 2009) have resulted in a vast accumulation of progressively higher quality phylogenies Maddison, 1997; e.g., Brassac & Blattner, 2015), leading to recognition of monophyletic groups and estimates of the temporal dynamics of evolutionary radiations (Nee, May, & Harvey, 1994; Magallón & Sanderson, 2001; Alfaro et al., 2009; Stadler, 2011; Morlon, 2014; Rabosky, 2014). These have revealed orders-of-magnitude differences in clade diversification rates, exemplified by *Amborella trichopoda*, an understory shrub endemic to New Caledonia, which is the only species of an angiosperm clade that is sister to, and therefore just as old as, the clade that contains all remaining ca. 400,000 species of flowering plants (Albert et al., 2013). Placing such salient diversification rate differences into a striking temporal context, Salzburger (2018, p. 705) recently noted that within “the time span that it took for 14 species of Darwin’s finches to evolve on the Galapagos archipelago […], about 1,000 cichlid species evolved in Lake Malawi alone”.

### Drivers of evolutionary radiations

The importance of **traits** and environments for understanding the mechanisms underlying evolutionary radiations was emphasized by Simpson (1944), and he postulated that most radiations are underpinned by **adaptation**. His adaptive radiation model envisioned diversification to take place in **adaptive zones**. A species may enter new adaptive zones by the evolution of new traits, climate change, or the formation of novel landscapes and disturbance regimes, such as newly emerged volcanic islands or human-induced night-light environments around the globe (Simpson, 1953; Erwin, 1992). The adaptive zone is a metaphor for the ways in which evolutionary innovations interact with environmental factors to modulate species **diversification** and trait **disparification** rates (de Vladar, Santos, & Szathmary, 2017; Olson, Arroyo-Santos, & Vergara-Silva, 2019), and Simpson’s concept of adaptive radiation underpinned and inspired a highly productive period of research on evolutionary radiations (e.g., Fryer, 1969; Losos, 1994; Sanderson & Donoghue, 1994; Baldwin, 1997; Hughes & Eastwood, 2006; Losos & Ricklefs, 2009; Wagner, Harmon, & Seehausen, 2012; Cooney et al., 2017; Marques, Meier, & Seehausen, 2019).

The effects of traits on species diversification (the interplay of speciation and extinction rates), combined with the temporal sequence of geographic movement and environmental change, estimated across a phylogenetic tree, has led to the recognition of **key innovations** (Miller, 1949; Van Valen, 1971; Liem, 1973; Sanderson & Donoghue, 1994; Heard & Hauser, 1995; Hunter, 1998). Key innovations are exemplified by freezing tolerance in Antarctic fishes (Portner, 2002) and herbaceous life-history strategies for occupying seasonally freezing environments by flowering plants (Zanne et al., 2014). In addition, phylogenetic comparative studies (Felsenstein, 1985; Harmon, 2018) have revealed the importance of **key events**, such as mass extinctions (e.g., of dinosaurs), climate change (e.g., late Miocene aridification), and orogeny (e.g., of the Andes and the New Zealand alps). Including evolutionary changes in genomic structure has led to the recognition that the connections between key innovations, key events, and diversification rate shifts can be complex (Erwin, 2001, 2017), for example, in context of hybridization and whole-genome duplications in flowering plants (Tank et al., 2015; Landis et al., 2018; Naciri & Linder, 2020) or African Rift Lake cichlids (Meier et al., 2017; Irisarri et al., 2018). Generally, however, it is the interaction between variable **intrinsic** (e.g., genome duplication) and **extrinsic** (e.g., climate change) factors that is thought to modulate diversification rates (for a review on context-dependent diversification see Donoghue & Sanderson, 2015; on the interplay of dispersal and biome shifts see Donoghue & Edwards, 2014).

The interaction between intrinsic (lineage-specific) and extrinsic (environmental) factors provides the **ecological opportunity** for adaptive radiations to occur (Erwin, 2015). Simpson (1953) summarised such opportunities in terms of three factors: (i) physical access to an environment, resulting from dispersal, or from the change of geo-ecological conditions in the region where a lineage already occurs; lack of effective competition in the environment, because no suitably adapted lineages already occur there; (iii) genetic capacity and adaptability of a lineage, which can be manifested in the evolution of key innovations, or more generally, in the ability to more readily explore the character space of certain trait innovations (Nürk, Atchison, & Hughes, 2019). All three conditions have to be met for successful adaptive radiation to start (Donoghue & Edwards, 2014; Stroud & Losos, 2016).

The evolution of diversity requires the evolution of reproductive isolation, which can be promoted by geographic fragmentation. Geographic isolation can result in stochastic divergence, i.e., by intensified genetic drift in smaller populations (Kimura, 1968; Duret, 2002), eventually leading to allopatric speciation. Repeated allopatric speciation can result in “non-adaptive” radiation (Gittenberger, 1991; Comes, Tribsch, & Bittkau, 2008; Verboom, Bergh, Haiden, Hoffmann, & Britton, 2015). Such (mainly) isolation-driven processes are contrasted to ecological speciation, which is driven by divergent selection pressures from the environment, implying that there will be (some) adaptation. Ecological speciation can result in repeated evolution of phenotypes and trait–environment interactions in adaptive radiations (trait utility; Schluter, 2000). However, in most radiations both ecological adaptation (natural selection) and geographic isolation (intensified genetic drift) are involved (Gittenberger, 2004; Brawand et al., 2014), although their relative contributions to diversification may vary between study systems (Rundell & Price, 2009; Czekanski-Moir & Rundell, 2019; Naciri & Linder, 2020).

Variation in the relative contributions of non-adaptive and adaptive processes to diversification was encapsulated by Simpson (1953) in his distinction between ‘access to an environment’ and ‘lack of competition in that environment’. In ecological niche theory, it is well appreciated that species live in environments that can be described by abiotic and biotic factors (for a review on the various aspects of niche concepts see McInerny & Etienne, 2012; McInerny, Etienne, & Higgins, 2012). Both abiotic and biotic factors can influence diversification rates (Holt, 2009) and may have varying or even opposite effects (Bailey, Dettman, Rainey, & Kassen, 2013) on speciation probability and extinction risk (Ezard, Aze, Pearson, & Purvis, 2011). Species interactions can drive diversification (Brodersen, Post, & Seehausen, 2018; Gavini, Ezcurra, & Aizen, 2019), or can have negative effects on species diversity, for example under competition for limited resources (Rosenblum et al., 2012; Harpole et al., 2016). On the other hand, heterogeneous abiotic conditions appear to be generally associated with higher species diversity (Rainey & Travisano, 1998; see also Erwin, 2001) potentially due to higher carrying capacities of larger and more heterogeneous areas (Field et al., 2009; Wagner, Harmon, & Seehausen, 2014; Storch & Okie, 2019), greater topographic complexity (Badgley et al., 2017; Sundaram et al., 2019), more climatic variability (Weigelt, Steinbauer, Cabral, & Kreft, 2016; Flantua, O’Dea, Onstein, Giraldo, & Hooghiemstra, 2019), or more complementary disturbance dynamics concordant or discordant with life histories (Jentsch & White, 2019). Indeed, abiotic and biotic factors and their effects on diversification are firmly established in research on evolutionary radiations (Ezard et al., 2011; Aguilée, Gascuel, Lambert, & Ferriere, 2018; Condamine, Romieu, & Guinot, 2019) and both factors are part of the extrinsic environment in which evolution of a lineage takes place.

In general, it is not the particular sequence of trait evolution or access to a novel environment that **triggers** a radiation, but the establishment/evolution of a complementary state – either the establishment of an environment that fits a pre-evolved trait or an **exaptation**, or the evolution of the trait that is an adaptation to a pre-existing environment (Kozak & Wiens, 2010; Wagner et al., 2012; Bouchenak-Khelladi, Onstein, Xing, Schwery, & Linder, 2015; Nürk, Michling, & Linder, 2018). Exploring the theoretical arguments that underpin such context-dependent radiations, Donoghue and Sanderson (2015) coined the terms **synnovation** for interacting combinations of (several) innovative traits, and **confluence** to describe sequential combinations of a set of traits and events along the stem lineages of radiating clades. The idea that evolutionary radiations are the product of synnovations and confluences of multiple intrinsic and extrinsic factors has gained momentum (Arakaki et al., 2011; Wagner et al., 2012; Guerrero, Rosas, Arroyo, & Wiens, 2013; Seehausen, 2015; Linder & Bouchenak-Khelladi, 2017; Harmon et al., 2019; Nürk, Atchison, et al., 2019).

## The Evolutionary Arena framework

### Description of the framework

Here we synthesize these insights into the drivers of evolutionary radiations – the context-dependent interplay between clade-specific intrinsic and extrinsic biotic and abiotic factors – into a simple framework, which we call the ‘Evolutionary Arena’ (EvA). In EvA the diversification (or disparification) rate of a focal lineage is a function of three components into which all macroevolution-relevant processes can be grouped and parameterised:

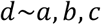

where *d* = **d**iversification or **d**isparification rate, *a* = **a**biotic environment, *b* = **b**iotic environment, and *c* = **c**lade-specific phenotypes or traits (Fig. 1).

**Figure 1.**
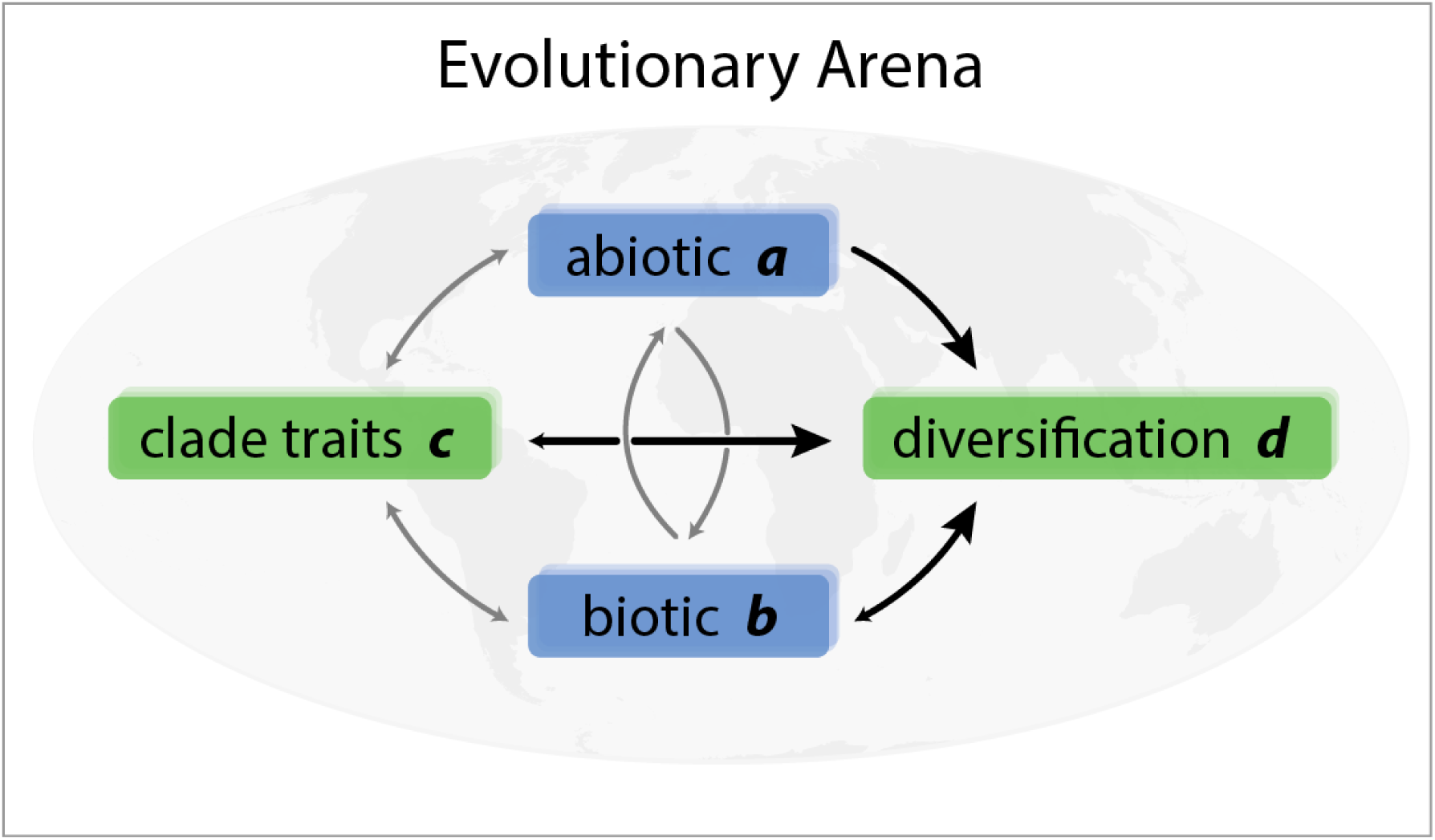
The Evolutionary Arena framework. The four components of the Evolutionary Arena (boxes) are illustrated with interactions (arrows) among the components, with those influencing diversification rates in black (larger arrows). We refer to the environment (abiotic [*a*] and the biotic [*b*] components; blue boxes) in combination with clade traits (*c*) and diversification / disparification rate (*d*) of an evolutionary lineage (green boxes) as the Evolutionary Arena. This framework is potentially dynamic, because interactions among the components allow for feedback, together shaping evolution. Note that, although all components can affect each other (positively or negatively; indicated by small and/or grey arrows), we focus here on the dependence of diversification on environmental (extrinsic biotic and abiotic) and clade-specific (intrinsic trait) factors.

In more detail, these four components are:

*d: diversification* or *disparification* is the rate of evolution in the broadest sense and depends on the values of the *a, b*, and *c* components (note, *d* might influence the other components as well; Fig. 1). Here, *d* can be expressed by the rate of change in taxonomic diversity (number of species), interspecific morphological/phenotypic disparity, DNA nucleotide diversity and genetic differentiation (e.g., functional variation of expressed genes or metabolites), physiological diversity (e.g., photosynthetic modes), or niche diversity (e.g., diversity of ecological niches occupied) and differentiation (e.g., rate of ecological expansion). This list is not exhaustive, and which expression of *d* is used depends on the question being investigated. The instantaneous rate of diversification, defined as speciation minus extinction per unit time (Nee et al., 1994), is the simplest expression of *d* and can be directly inferred from a dated species-level phylogeny (Magallón & Sanderson, 2001). Disparification can be measured, for example, by (relative) evolutionary rate estimates (Butler & King, 2004), the change in trait variance in a clade through time (Rolshausen, Davies, & Hendry, 2018), or by transition rates between discrete character states (Huelsenbeck, Nielsen, & Bollback, 2003). The diversification rate may be positive, resulting in an increase in diversity, or negative, resulting in a decrease in diversity.

*a: abiotic environment* incorporates abiotic factors, such as climate, soil, habitat variables, or fragmentation of the species range. It can also include biotic elements, particularly with respect to their physical characteristics, such as vegetation types (which, as classes, cannot evolve). Component *a* can be measured as absolute values, e.g., area or niche space, or as the variance in these across space and time, e.g., variance in mean annual precipitation, in number of vegetation types or soil types, physiographic heterogeneity, and the functional and structural connectivity. Ideally, the abiotic environment is described by processes that generate this environment, such as erosion or orogeny, or patterns of change, such as climate or vegetation change.

*b: biotic environment* captures the interactions of the focal lineage with all other species (including species both within and outside the clade and also including trophic interactions). The interaction(s) can be, for example, mutualistic or commensalistic (e.g. pollinators or dispersers, interspecific facilitation, or mycorrhiza), antagonistic (e.g., herbivores, diseases, interspecific competition, or hosts for parasites), or genetic (e.g., hybridization/introgression and horizontal gene transfer). Note, however, that the *capacity* to hybridize may be treated as a trait, and so categorised under *c*. These biotic interactions can also be indirect, if seen as part of the extended phenotype of the focal lineage (e.g., habitat modification such as niche construction, or grasslands increasing fire frequency, or stoichiometric needs of organisms modifying available resources; Jentsch & White, 2019). There is a rich theory surrounding biotic interactions (e.g., the macro-evolutionary Red Queen hypothesis [Benton, 2009, see also Van Valen, 1973], niche construction [Laland, Matthews, & Feldman, 2016]) suggesting they can have a powerful influence on diversification rates. However, the biotic environment is complex; quantifying and including it in comparative analyses is a challenging task (Harmon et al., 2019). Indeed, recent studies indicate that inter-lineage competition (e.g., Silvestro, Antonelli, Salamin, & Quental, 2015; Pires, Silvestro, & Quental, 2017) or interactions with possible dispersal agents (Onstein et al., 2018) may modulate diversification rates.

*c: clade-specific traits* include the phenotypic characteristics of the focal species or lineage. Included here are, among others, physiological characteristics (e.g., variation in photosynthetic mode), anatomical or morphological traits, life history strategies, pollination strategies and dispersal modes. Clade-specific traits are part of the phenotype and can be labile or phylogenetically conserved. This is illustrated by the remarkable floral variation in the orchid genus *Disa* (Johnson, Linder, & Steiner, 1998), and the impact of a lack of floral morphology variation on diversification in the oil-bee pollinated Malpighiaceae (Davis et al., 2014). The phenotype (or the extended phenotype) is a manifestation, or function, of the genome (via the genotype–phenotype map, inasmuch as this is independent of the environment), or genome diversity (e.g., structural variation such as ploidy level variation, variation in genome size, or DNA nucleotide sequence variation) within the focal group. Therefore, this parameter should ultimately be genetically measurable.

The formulation of the EvA framework is general because it is simple and all-encompassing. The key challenge faced in studies addressing evolutionary radiations is to disentangle the effects of the different components at different moments in time. The ultimate in understanding radiations and evolutionary stasis would be the joint estimation of all components at all times, but we cannot analyse an infinite number of variables. On the other hand, studies sometimes assign an increase or decrease of diversification to particular factors that happen to have been investigated and quantified, disregarding the possible effects of other factors that may be driving or constraining diversification. We argue that the complete EvA, as expressed by the abiotic and biotic environment, as well as the traits of organisms, should be considered when attempting to explain variation in diversification rates. Not only does this ensure that all relevant factors are taken into account, but the same set of components are considered in all analyses. In this way, the EvA framework can provide new insights by comparing diversification between clades directly.

To make EvA operational requires parameterising it appropriately, which means making it more specific and detailed. Here, we describe four simple extensions to illustrate how the EvA framework can be enriched by more properties to provide insights into different hypotheses in evolutionary diversification. We end this section by outlining general analytical approaches and possibilities for null hypothesis formulation.

1. *Direction of effects*: The three components *a, b*, and *c* can have a significant positive (+) or negative (-) effect on *d*, thus causing the diversification rate to increase or decrease, or even show a false absence of change as the summed end result of the three. This process of ‘nullification’, or less increase or decrease than expected, is sometimes ignored and interpreted as a lack of power by the factors influencing diversification. Statistical approaches comparing different systems could provide insights in cases in which diversification rates are higher or lower than expected.
2. *Complex conditions*: The framework can be expanded to include any set of variables per component. For example, the abiotic environment could be described by climatic factors such as mean annual precipitation, temperature, and by disturbance factors such as fire frequency. The set of variables selected depends on the hypothesis being tested. If, for example, the hypothesis is that *d* is related to the total variance in the abiotic environment *a*, then 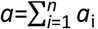, where *a*_i_ is the *i*^th^ variable of *a* (up to the last variable *a*_*n*_) given as a measure of variance. Thus, the abiotic environment *a* (as well as the other components in EvA) can be decomposed into diverse measures.
3. *Rate shifts*: Explanations for shifts in diversification rates can be sought by testing for changes in the EvA component values. This can be done by including initial values at time *t* (ancestral) and the values after a time interval Δ*t* (derived), for example, at time *t*+Δ*t* after an event: Δ*d* = (*a, b, c*)_*t*+Δ*t*_ − (*a, b, c*)_*t*_ or simply Δ*d*∼Δ*a*, Δ*b*, Δ*c*. If we, for example, hypothesise that geographic movement to a new region is a key event, the condition of *a* varies between *t* and *t*+Δ*t*.
4. *Interaction of components*. We can incorporate interactions between components, such as *ac, bc*, or *abc*: *d*∼*a, b, c, ab, ac, bc, abc*. This allows us to analyse context-dependence: whether the single components, or the interactions among them modulate the diversification rate, i.e., driving or constraining the evolution of diversity. Such interactions can be exemplified by ecosystem engineers modifying disturbance regimes (Jentsch & White, 2019), such as impacts of grass invasion on forests increasing fire frequency and ultimately transforming the environment (Bond, Woodward, & Midgley, 2005; Beerling & Osborne, 2006). Similar interactions (or transformations) have been proposed for changes in fruit size (component *c*) as consequences of mega-herbivore extinctions (component *b*) and climate change (component *a*; Onstein et al., 2018). A further dynamic factor is provided by interactions between the diversification rate (*d*) and the predictor variables (*a, b, c*). Such feedback mechanisms can broadly be summarised by the concept of niche construction (Laland et al., 2016), and are exemplified by the ‘Viking Syndrome’ described in reference to grasses (Linder, Lehmann, Archibald, Osborne, & Richardson, 2018). The latter proposes that global grass success (in species richness, environmental range, ecological dominance and geographical distribution) is due to the high invasiveness of grasses, which results from their high rate of dispersal, effective establishment, ecological flexibility and disturbance tolerance (all component *c*), and ability to transform environments by increasing the frequency of fire (component *a*) and the density of grazers (component *b*). These scenarios describe feedback systems where through increased diversity and dominance, the diversifying clade increasingly modifies the abiotic and biotic environment.

In this way, the EvA framework can facilitate the direct testing of competing hypotheses about the diversification of a group, using standard model selection approaches (e.g., likelihood ratio tests for nested models, AIC or Bayes factors; Burnham & Anderson, 2002). Phylogenetic pseudoreplication (Maddison & FitzJohn, 2015), which describes the non-independence of, or the autocorrelation among, species’ traits due to shared ancestry, is a basic property of comparative analyses (Felsenstein, 1985). Phylogenetically independent contrasts (PIC; Felsenstein, 1985) and phylogenetic generalised least squares (PGLS; Grafen, 1989; Martins & Hansen, 1997) are methods for analysing comparative phylogenetic data by accounting for the covariances between traits resulting from shared phylogenetic history. These methods may be generally useful for exploring the relationships between components in the EvA framework. Although both methods can be thought of as “analogous to data transformations made to better approximate the assumptions of standard statistical tests” (Huey, Garland, & Turelli, 2019, p 762), they, as well as most other phylogenetic comparative methods (but see Rolshausen et al., 2018), implicitly assume a specific process underlying character state change along the evolutionary lineage (i.e., a model of character evolution such as Brownian motion; Boucher, Demery, Conti, Harmon, & Uyeda, 2018). The appropriate data transformation model in relation to the hypothesis being tested is a non-trivial question in comparative analyses (Uyeda, Zenil-Ferguson, & Pennell, 2018); graphical models depicting hypothesized causal links (Höhna et al., 2014) can help here. On the other hand, the case of a singular, unreplicated event in the evolutionary history of a lineage challenges the statistical power of comparative phylogenetic methods (Maddison & FitzJohn, 2015). An approach to overcome this limitation might be the combined application of hypothesis testing and exploratory methods (‘phylogenetic natural history’) as outlined by Uyeda et al. (2018).

Associations between diversification rates and the other components in the EvA framework can be tested using state-dependent speciation and extinction (SSE) models (such as BiSSE): a birth–death process where the diversification rates depend on the state of an evolving (binary) character (Maddison, Midford, & Otto, 2007), given a phylogeny and trait data. For complex components, the multistate SSE (MuSSE) model can be applied to accommodate several qualitative character states, or multiple (binary state) characters following Pagel (1994) for state recoding (FitzJohn, 2012). Although SSE model extensions for quantitative characters (QuaSSE; (FitzJohn, 2010) and geographic range evolution (GeoSSE; Goldberg, Lancaster, & Ree, 2011) (to name just a few; for a critical introduction to SSE models see O’Meara & Beaulieu, 2016) have been developed, in the SSE model-family none is currently available for the likelihood calculation of a model considering both quantitative and qualitative variables (but see Felsenstein, 2012; Revell, 2014). Nor do current SSE models allow the analysis of interactions, and so context-dependent radiations. This situation could be common under the EvA framework and represents a priority for method development.

Comparing a biologically meaningful and appropriately complex null hypothesis to the goodness-of-fit of alternative (H1) models is essential for detecting whether a character state-dependent model can explain more of the observed variation than could be expected under random diversification rates (Caetano, O’Meara, & Beaulieu, 2018). It has, for example, been shown for SSE models that the variation in the diversification rate observed in a phylogenetic tree is not necessarily explained by the focal factor (character) under study (Rabosky & Goldberg, 2015). False-positives can potentially result because the null hypotheses did not account for the possibility that diversification rates can be “independent of the character but not constant through time” (Harmon, 2018, p 215). Hidden state model (HSM) approaches (Marazzi et al., 2012; Beaulieu, O’Meara, & Donoghue, 2013; Beaulieu & O’Meara, 2016; Caetano et al., 2018), which incorporate unobserved (‘hidden’) factors as model parameters equivalent to observed ones, offer a solution to this problem. Comparing goodness-of-fit between ‘hidden state’ null models and those representing the focal factor(s) provides appropriately complex null hypotheses that can be used for testing differently parameterised EvA models, and thereby allows identification of “the meaningful impact of [the] suspected ‘driver[s]’ of diversification” (Caetano et al., 2018, p 2308).

### Advantages of the Evolutionary Arena framework

The Evolutionary Arena framework does not contribute any new concepts or terms but is built on the concepts developed over the past few decades, reviewed above (section “Drivers of evolutionary radiations”). However, what is still lacking is, as noted by Donoghue and Sanderson (2015), a single, simple formula with which to develop convincing hypotheses of the drivers of evolutionary rate changes. This we attempt to provide with EvA. The basic four components – diversification / disparification, clade-specific intrinsic traits, extrinsic abiotic and biotic factors – and their interactions can be compared between systems to gain more general insights into the factors that underpin evolutionary diversification. Considering all four components together in a single framework fosters a holistic approach. EvA consequently incorporates the full complexity of triggers, synnovations, and confluences associated with evolutionary radiations (Bouchenak-Khelladi et al., 2015; Donoghue & Sanderson, 2015). This facilitates comparative analyses of evolutionary radiations, or evolutionary stasis and decline, using phylogenetic comparative methods. This is possible because *d* can be positive or negative/smaller, so the correlates (e.g., *a* x *c* in EvA) of diversification increase or decrease (e.g., density-dependent slowdowns) can be sought. EvA does not present any new analytical methods, analyses within this framework can be done using existing packages and software (it may also indicate priorities for method development). Particularly important is the central notion that no single factor is a sufficient explanation for an evolutionary rate change, but that the interaction between external and internal factors results in shifts in diversification and/or disparification rates (Givnish, 2015). Overall, there are three heuristic advantages to couching evolutionary radiation studies in the EvA framework:

Firstly, this framework, similar to a model, predicts which factors may be drivers of evolutionary radiations. This reduces the risk of missing important drivers, and so stimulates the development of comprehensive models, rather the simple exploration of the effect of a factor on diversification rates. In addition, it encourages taking recent advances in understanding context-dependence into account.

Secondly, this framework is readily quantified, for example as a regression model. Quantification both facilitates and encourages data transparency (i.e. what data sets are used, and how these data are transformed). This transparency becomes more important as the model is expanded to reflect the complexity of the predictor factors.

Thirdly, it provides a single, general framework within which to analyse all or any evolutionary radiations. The framework can be applied to any biological organisms, geographical regions, or ecosystems. This facilitates the comparison among taxa and regions as to the processes underlying diversification, even if the studies were by different people. This will ease the progression from case studies to general syntheses.

### Case study: Conifers

We use a case study of conifer radiation to illustrate EvA implementation and component quantification. In contrast to the conceptual simplicity of the framework, obtaining data for all components across multiple lineages can be challenging, although well-developed phylogenies over a wide range of taxa are increasingly available. Here, we use published data on 455 conifer species (Larcombe, Jordan, Bryant, & Higgins, 2018) that enable parameterisation of the *d*∼*a, b, c* framework. The conifers provide an excellent study clade for comparative analysis: the lineage is rich in species grouped into well-defined clades, geographically widespread, and well-studied with excellent distribution data (Farjon, 2018). Although conifers originated ca. 300 million years (Ma) ago, with the main clades thought to have diverged between the early Triassic (ca. 240 Ma) and mid-Cretaceous (ca. 100 Ma), most modern species arose in the Neogene or Quaternary (23 Ma–present; Leslie et al., 2012). We used the dated phylogeny of Leslie et al. (2012; inferred from a Bayesian analysis assuming an uncorrelated lognormal clock model, based on two nuclear and two plastid genes, and calibrated with 16 fossils) to define 70 reciprocally monophyletic or single-species groups; using a stem age cut-off at 33.9 Ma (the Eocene/Oligocene boundary) in order to focus on the variables which could explain Neogene–Quaternary diversification rate variation (Larcombe et al., 2018). Forty-one of these groups have more than one species, the most species-rich 52 species, and range in age from 34 to 146 Ma.

The factors that contribute to *a, b* and *c* in the conifers model were derived from the output of a process-based niche model (Larcombe et al., 2018). This niche modelling method is described in detail in Higgins et al. (2012), and is based on a mechanistic model of plant growth, the ‘Thornley Transport Resistance’ model, which models resource acquisition, transport and allocation between roots and shoots, based on environmental information extracted from species distribution data. This produces two types of information for each species: 1) estimates of the geographic distribution of the potential niche of each species (i.e., a species distribution model [SDM]); and 2) estimates of the physiological parameters that describe the niche of each species (Higgins et al., 2012). We used these metrics, and occurrence data and species richness per clade to parametrise *d*∼*a, b, c* as follows. Note that all four factors are values for the 41 multispecies clades, not for the constituent species individually (Larcombe et al., 2018):

*d* = the diversification rates for each clade: calculated using the method-of-moments estimator of Magallón and Sanderson (2001) as 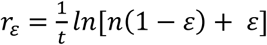, where *r*_*ε*_ is the net diversification rate assuming a relative extinction fraction *ε* = 0.9, *n* the number of extant species, and *t* the stem age of the clade.

*a* = abiotic environment, quantified by the clade’s potentially suitable area size: the projected geographical range reflecting the potential niche of all species within the clades. This is calculated per clade as the number of ¼° grid cells across the globe that at least one species of the clade can occupy, based on the physiological SDM and corrected for clade species numbers (i.e., rarefied to the clade with the lowest diversity to remove sampling effects; Larcombe et al., 2018). This means that if the score is small, the niche size of a clade is expected to be narrow, and if the score is large, the niche size of the clade is large so that the clade comprises ecologically more generalist species or the species in the clade might be specialised but different from one another. Despite its simplifying assumptions about the spatial distribution of environmental variation (some types of environment are more common than others), this clade-wise suitable area size is an appealing measure of *a* because it approximates the potential niche, consequently biotic interactions and effects of traits can be estimated separately. Figure 3 shows the combined potential niche for all 455 species in the dataset, i.e., the abiotic arena of the conifers.

**Figure 2.**
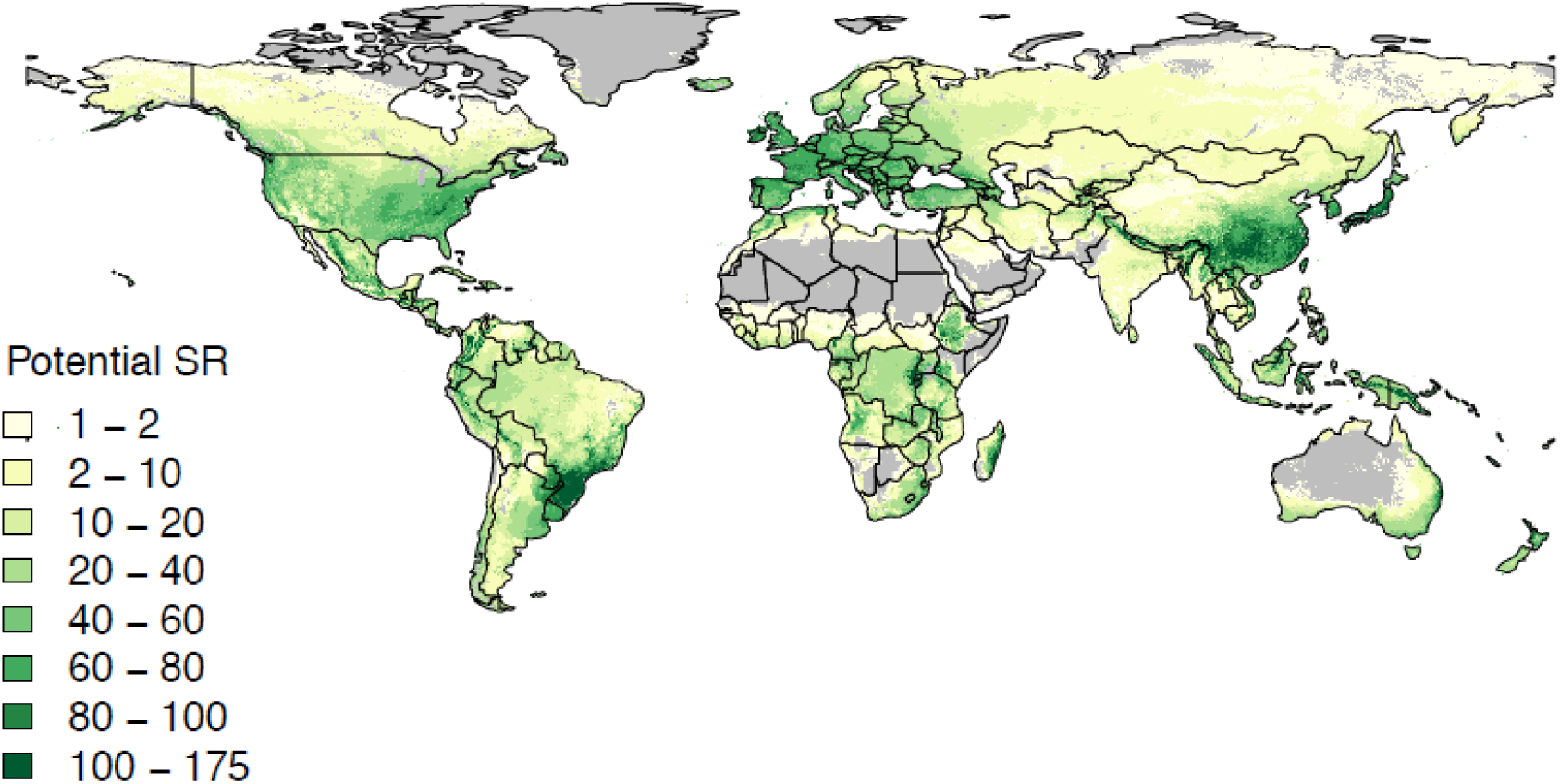
Projected potential species richness (SR) of conifers representing their abiotic arena. The abiotic arena of conifers is defined by geographic locations (quarter degree grid cells) that can support one or more of the 455 conifer species, based on projections from the process-based physiological niche model (approximating the fundamental conifer niche). The projected potential SR (yellow-green shading) highlights areas that are suitable for conifer species. The abiotic arena alone does not always predict patterns of conifer diversity. For example, the eastern Congo is predicted to be climatically suitable for many species but has relatively low conifer diversity (Farjon, 2018). It is likely that clade-specific traits (*c*) and biotic interactions (*b*) limit the diversity in certain regions (see main text).

**Figure 3.**
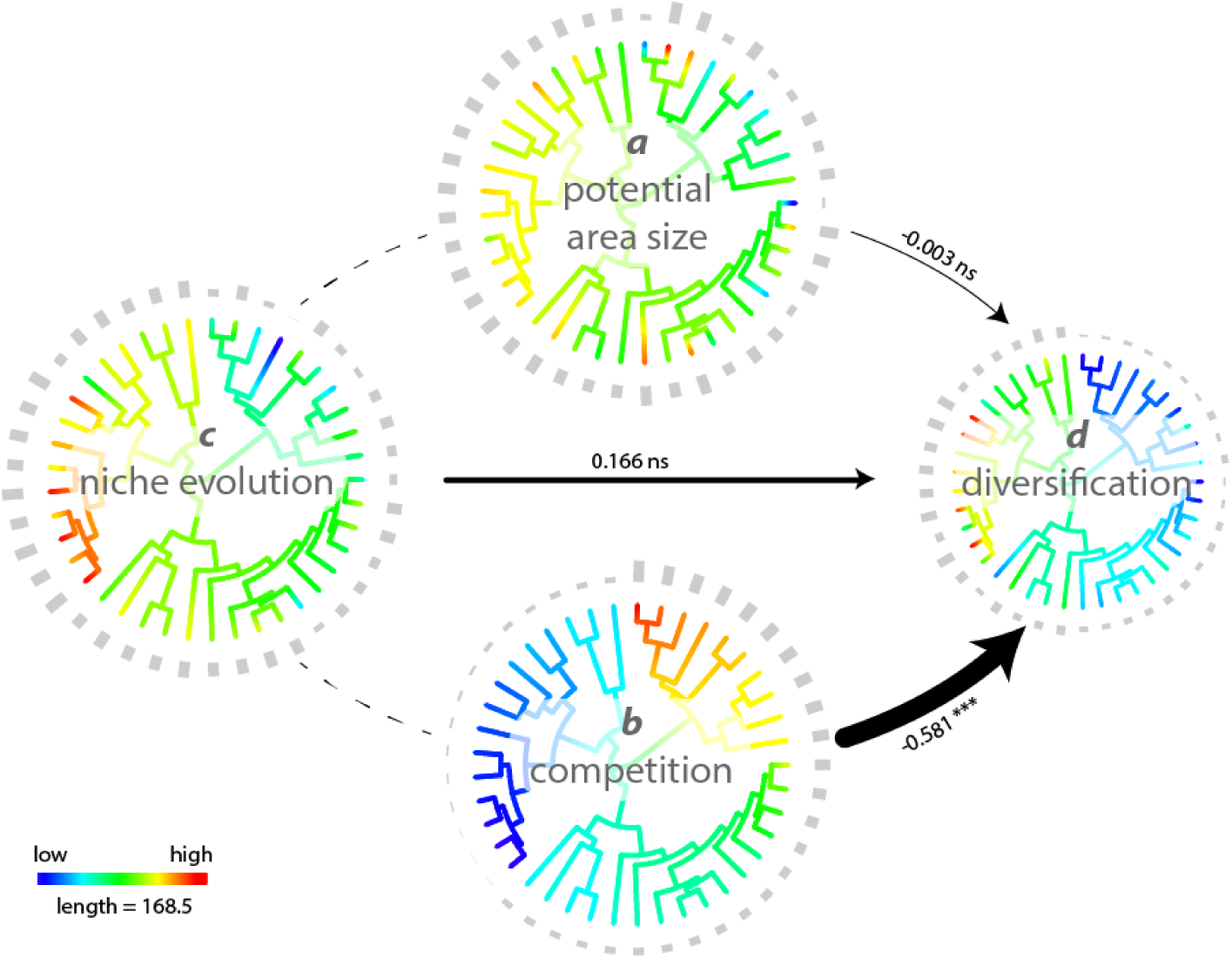
Quantified EvA model for the conifers. Coloured trees illustrate the distribution of abiotic environment (*a*), competitive interactions (*b*), rate of niche evolution (*c*), and diversification rates (*d*) across the conifer phylogeny with bars at tips detailing the values per clade and variable. The estimated effects on net diversification rates *r*_*ε*_ are indicated by arrows scaled to the standardised coefficients (slopes) also showing significant levels (ns, non-significant; ***, P <0.001). Colours on branches are rate estimates (obtained using the R package phytools’ fastAnc function; Revell, 2012; see R scripts in Nürk, Linder, et al., 2019) and illustrate parameter distribution on the tree. When competition among species is low and the rate of niche evolution in a clade is pronounced, the diversification rate of that clade accelerates.

*b* = competitive interactions estimated at the species level: we determine the expected competition for each species with all members within its clade (one of the 41 clades defined above) as the product of niche and geographic overlap between species. The metric underestimates competition because it excludes competitive processes with other clades and other indirect competitive processes (see discussion in Larcombe et al., 2018). Geographic overlap is estimated based on the occurrence data. Niche overlap between each species pair was calculated using Schoener’s niche overlap metric *D* (Schoener, 1968) based on the potential distributions from the SDM analysis. We then scaled these two numbers to range from 0 to 1 for each species pair and multiplied them to provide a competition index (Larcombe et al., 2018). This means that if either score is zero, the competition score is zero, and if they have the same (potential) niche and the same (realized) range then the competition score is 1. The species-level estimates were averaged to provide a clade-level competition score.

*c* = clade-specific rate of niche evolution: we used eleven physiological traits (see Fig. 6 in Larcombe et al., 2018) that were identified as being most important for defining the overall niche space of conifers (Larcombe et al., 2018). Although an effectively limitless number of physiological traits could be defined, our method provides an objective selection criterion of ecologically appropriate measures. These eleven traits were fitted together in a multivariate Brownian motion model of evolution (Butler & King, 2004) on the conifer phylogeny of Leslie et al. (2012), and the diagonal elements of the resulting variance–covariance matrix for the species traits represent the phylogenetic rate of evolution (O’Meara, Ané, Sanderson, & Wainwright, 2006). These were summed and scaled to provide a multidimensional clade-level niche evolution rate.

**Figure 4.**
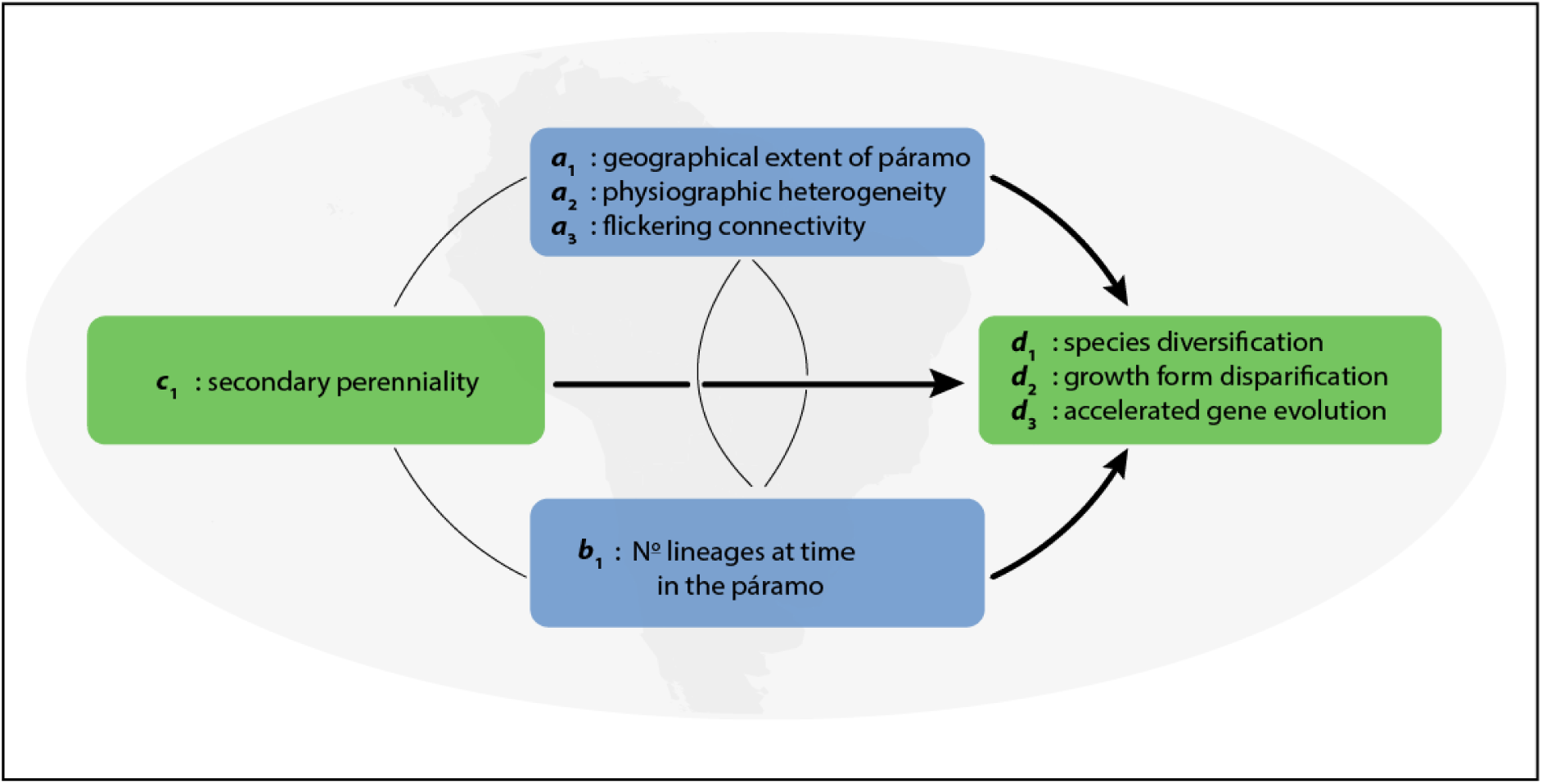
The EvA model for *Lupinus*: (*d*_1_, *d*_2_, *d*_3_) ∼ (*a*_1_, *a*_2_, *a*_3_), *b*_1_, *c*_1_. Note that rates of growth form disparification and accelerated gene evolution can influence diversification rates; consequently, the grouping of variables as the components of the EvA framework depends on the questions being investigated.

**Figure 5.**
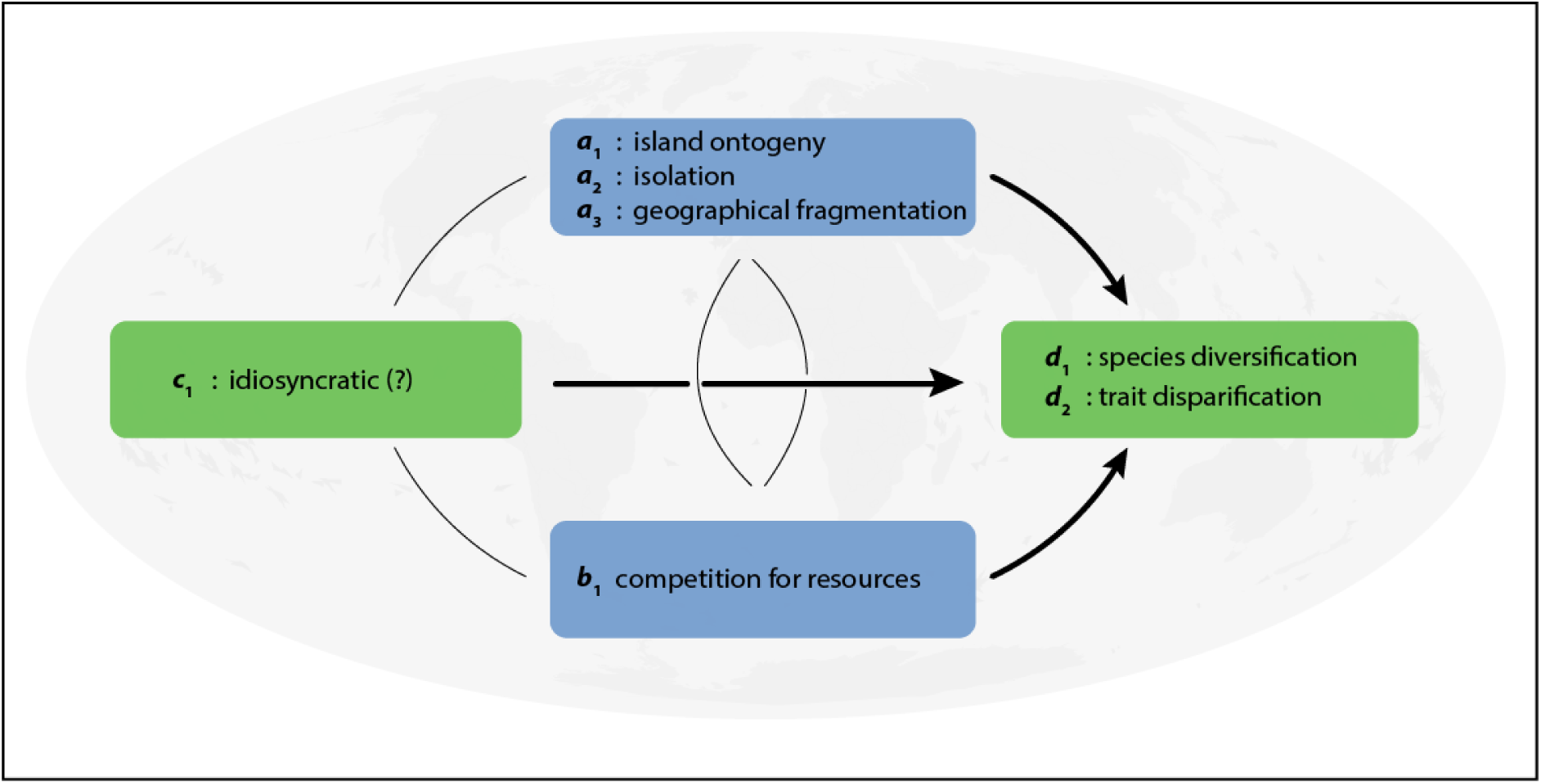
The EvA model for island radiations: (*d*_1,_ *d*_2_) ∼ (*a*_1_, *a*_2_, *a*_3_), *b*_1_, *c*_1_. Island ontogeny describes the typical life-cycle of oceanic islands; isolation the degree of isolation of the insular system (e.g. distance from the continent); geographical fragmentation could be the number of islands (if dealing with an archipelago). Incorporating the lineage-specific traits is complex, as several independent lineages may be involved. Trait disparification is particularly interesting in islands, for example, Hawaiian honeycreepers and silverswords.

**Figure 6.**
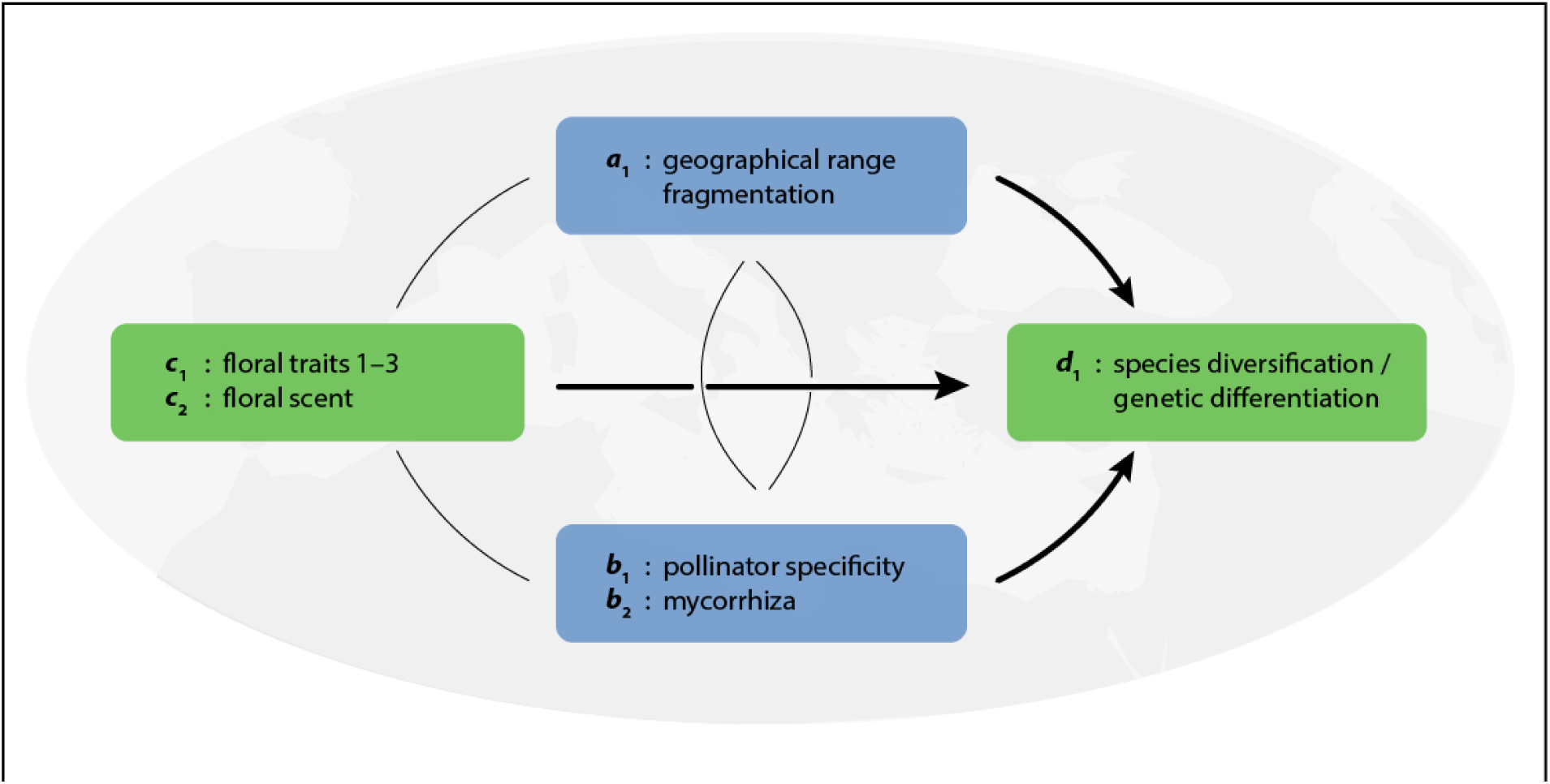
The EvA model for *Ophrys*: *d*_1_ ∼ *a*_1_, *b*_1_, (*c*_1_, *c*_2_). Floral traits (*c*_1.1_ geometry, *c*_1.2_ epidermis micromorphology, *c*_1.3_ colour and patterns), also including scent, are probably the most important regulators of the highly specific pollination system, but the role of mycorrhiza and geographical isolation on the Mediterranean islands is poorly understood.

Our expectation is that conifer diversification rates (*d*) are positively affected by the available abiotic environment (*a*) and the rate of niche evolution (*c*), and negatively by inter-specific competition (*b*). We fitted the conifers EvA model 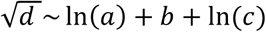 by means of phylogenetic generalised least squares (PGLS), controlling for the non-independence between cases resulting from phylogenetic structure in the data using the R v3.5.3 (R Core Team, 2013) library ‘phylolm’ v2.6 (Ho & Ané, 2014). Note that the variables *a, c* and *d* were transformed to reduce skewness in the data. PGLS estimates the regression parameters of all variables (the scaled variables used to parameterise the components of EvA), adjusted for the phylogenetic signal in the model residuals. We accessed the standardised coefficients and calculated the variance explained by the full model using the coefficient of determination (R^2^) to measure goodness-of-fit, and also assessing partial r^2^ (variance explained per predictor variable *a, b*, and *c*) using the R library ‘rr2’ v1.0.1 (Ives & Li, 2018; Ives, 2019). R scripts are available in Dryad (Nürk, Linder, et al., 2019).

The full model accounted for 64% of variation in diversification rates among the conifer clades (adjusted R^2^ = 0.638). The predictor variables *a, b*, and *c* in the conifer EvA model differentially contributed to explaining *d* (table 1). Against our expectations, the abiotic environment of clades (*a*) showed no relationship to diversification rate (slope -0.003, P = 0.89), neither did the rate of niche evolution (*c*; slope 0.166, P = 0.36). Contrarily, competitive interactions (*b*) between the species in a clade indicated a significant negative relationship to diversification (slope -0.581, P < 0.001), supporting our expectation.

**Table 1.** Estimated effects of abiotic environment (*a*), competitive interactions (*b*), and rate of niche evolution (*c*), on net diversification rate r_ε_ (*d*) across the conifer clades (samples size n = 41).

| Source | Slope | Std. error | t value | P | partial *r*<sup>2</sup> |
| --- | --- | --- | --- | --- | --- |
| Abiotic environment (*a*) | -0.003 | 0.148 | -0.018 | 0.985 | 0.000 |
| Competition (*b*) | -0.581 | 0.127 | 4.563 | <0.001 | 0.360 |
| Rate of niche evolution (*c*) | 0.166 | 0.181 | 0.919 | 0.364 | 0.022 |
| (Intercept) | 0.601 | 0.160 | 3.761 | <0.001 | — |

The significantly negative effect of competition (*b*) on diversification (*d*) indicates higher diversification rates in clades where competition among species is low. This result is consistent with the concept of diversity-dependent diversification (Foote, 2000; Rabosky, 2013) and suggests that diversity-dependent relationships are more important among the conifers in regulating diversification rate than potential area size (*a*) or rates of physiological trait evolution (niche evolution; *c*). However, the rates of niche evolution (*c*) among the conifer clades show a very similar, although inverse, pattern to that of competition (*b*) (Fig. 4). Estimates of a model accounting for interactions among predictor variables (results not shown due to lack of statistical power using n = 41; see R scripts available in Nürk, Linder, et al., 2019) indicated that the two-way interaction *b*:*c* (competition interacting with rate of niche evolution) influenced diversification in the conifers, in line with findings by Larcombe et al. (2018), who showed that conifer evolution is jointly shaped by bounded and unbounded evolutionary processes (e.g., Harmon & Harrison, 2015). The two-way interaction *b:c* may enhance or relax diversity-dependent processes so as to promote or constrain diversification (Larcombe et al., 2018). This is also consistent with the concept that spatial (or temporal) variation in trait disparity can result in variation in competitive pressure (McPeek, 2008; Marshall & Quental, 2016). The interaction of competition and rate of niche evolution suggests that the fastest diversification in conifers is found when competition is low (increased ecological opportunity), which could be the result of fast trait/niche evolution (high adaptability of the lineage).

The case presented here shows the potential to infer general patterns using the EvA framework. However, it is in no way a full exploration of the approach, and more sophisticated analyses are likely to prove more informative. For example, our analysis assumes that rates for *d, b*, and *c* are fixed, which is an over-simplification. It could be interesting to repeat the analysis using species instead of clades, as this allows us to account for phylogenetic structure within the clades, ecologically highly variable species, and diversification stasis. However, there are issues interpreting tip-diversification rates (Title & Rabosky, 2019). Methods are available to reconstruct ancestral sympatry and infer the effect of competition on trait divergence and lineage diversification (Aristide & Morlon, 2019; Harmon et al., 2019). Methods that reconstruct and evolve ancestral states along phylogenies for *a, b*, and *c*, are also appealing (Uyeda et al., 2018), and with increasing complexity of the EvA model, approaches such as hidden states will be important for rigorous testing against equivalently complex null hypotheses (Caetano et al., 2018). This illustrates the value of EvA in making data assumptions explicit.

### Conceptual examples

#### *Lupinus* continental radiation

Among the most intensively investigated radiations are several in the tropical alpine environments of the high-elevation Andean grasslands. These environments emerged as a result of the most recent Pliocene uplift of the Northern Andes, and consequently the radiations themselves are largely confined to the Pleistocene (Luebert & Weigend, 2014; Hughes & Atchison, 2015). These are exemplified by the diversification of *c*. 85 species of *Lupinus* L. (lupines, atmospheric nitrogen-fixing Leguminosae) within the last 1.2–3.5 Ma. The Andean *Lupinus* radiation has been attributed to a combination of intrinsic evolutionary (trait) innovation and extrinsic ecological opportunity (Hughes & Atchison, 2015). The shift from an annual to a perennial life history (i.e., evolution of a clade-specific phenotype = *c* in EvA; Fig. 4) is hypothesised to have acted as a key innovation facilitating occupation of mesic montane habitats (Drummond, Eastwood, Miotto, & Hughes, 2012), also enabling accelerated disparification of plant growth forms in the Andes (Nürk, Atchison, et al., 2019). This is because perennials have different cold tolerance strategies than annuals, underpinning their adaptation to high-elevation ecosystems, and a fundamentally greater potential growth-form disparity than annuals (Ogburn & Edwards, 2015; Nürk, Atchison, et al., 2019). At the same time, extrinsic ecological opportunities for diversification were available in the island-like high-elevation habitats that emerged during the last few million years due to Andean uplift and cooling of global temperatures (i.e., abiotic factors = *a* in EvA), prompting Hughes and Eastwood (2006) to refer to the Andean *Lupinus* clade as an example of “island-like radiation on a continental scale”. In this example, the evolution of secondary perenniality, the shift from lowland to montane habitats, and the primary shift to higher rates of species diversification all coincide on the same branch of the phylogeny, presenting an example of a “key confluence” *sensu* Donoghue and Sanderson (2015). In the Andean *Lupinus* clade, *d* has been estimated as the rate of species diversification (Drummond et al., 2012), phenotypic trait evolution or disparification of plant growth forms (Nürk, Atchison, et al., 2019), and coding DNA sequence evolution (Nevado, Atchison, Hughes, & Filatov, 2016). All of these estimates of *d* show accelerated rates across the western New World montane radiation when compared to the earlier diverging lineages of more slowly diversifying lowland western New World *Lupinus* annuals.

*Lupinus* thus provides an apparently straightforward example of a confluence of intrinsic innovation and extrinsic opportunity as the trigger for accelerated diversification of species and disparification of plant growth forms. This explanation assumes that ecological opportunity presented by empty sky island habitats and the means to take advantage of those opportunities (secondary perenniality) are driving diversification. However, treating the biotic environment (*b* in EvA) as zero, or empty of competition is clearly an over-simplification, given that there were apparently many plant radiations playing out during the Pleistocene across the high-elevation Andean grasslands, presumably in parallel with each other (Luebert & Weigend, 2014). The detailed order of timing of these radiations and their interactions remain unknown, just as interactions among sympatric and more or less contemporaneous radiations have been difficult to tease out more generally (Tanentzap et al., 2015). The EvA framework draws attention to the fact that biotic factors have not been critically investigated beyond the simple idea of lack of competition in the newly emerged tropical alpine sky island habitat (Fig. 5).

A more detailed analysis might indicate what factors of these high-elevation habitats are important for the observed high rates of diversification. Indeed, it is aspects of the abiotic environment (*a* in EvA) that are most often put forward as the central explanation for the numerous rapid recent radiations in the high-elevation Andean grassland. Foremost amongst these aspects are: (i) the large continental-scale extent of the high-elevation Andes; (ii) the extreme physiographic heterogeneity; and (iii) the rapid fluctuation in the extent and connectivity between the north Andean alpine sky islands (páramo) during the Pleistocene glacial-interglacial climate cycles. Physiographic heterogeneity of the Andes, spanning steep and extended environmental gradients (e.g., temperature and rainfall), has long been considered as a key factor driving Andean radiations (Hughes & Eastwood, 2006) and indeed of diversification more generally (e.g., Rangel et al., 2018). It has also long been recognised that the area and connectivity of the high-elevation Andean grasslands have varied dramatically through the Pleistocene due to elevational shifts in vegetation zones and species distributions imposed by glacial–interglacial periods. However, it is only recently that area and connectivity have been modelled and quantified in sufficient detail through the Pleistocene (Flantua et al., 2019) to assess the potential of such an alpine ‘flickering connectivity system’ (Flantua & Hooghiemstra, 2018) to further enhance diversification (e.g., Nevado et al., 2016). Such models demonstrate the need to quantify attributes of the abiotic environment (*a* in EvA) through time as well as the potential of such time-dependent models to make more realistic estimates of the impact on diversification rates (*d* in EvA).

Considering the Andean *Lupinus* radiation in the light of the EvA framework highlights the lack of knowledge of the biotic interactions involved (*b*). This illuminates that explanations using only the abiotic environment (*a*) and intrinsic traits (*c*) may be incomplete. Using the EvA framework suggests new research questions.

#### Island radiations

Volcanic islands that arise *de novo* on the oceanic crust show a typical lifecycle. In contrast to islands on the continental shelf, oceanic islands emerge following a volcanic eruption, grow rapidly in area and elevation, and then erode down to the sea level over 5 to 30 million years, depending on the substrate and climatic conditions. Even though oceanic islands may show a flickering connectivity effect as a result of Pleistocene climate fluctuations (e.g., lower sea levels would have resulted in larger islands areas leaving an imprint on current biodiversity; Weigelt et al., 2016), we here focus on the entire oceanic island lifecycle (Whittaker, Triantis, & Ladle, 2008; Borregaard et al., 2017). At this scale, island ontogeny can be considered unimodal in its key properties, and so differs from a flickering model as described for terrestrial high mountains (Flantua & Hooghiemstra, 2018; Flantua et al., 2019), which is multimodal. The island lifecycle is an ontogenetic geomorphological trajectory of area, elevation and habitat diversity – from island birth, through maturity, until island submergence. Consequently, this can be analysed as a continuous time series. The potential effects of this ontogeny on evolutionary processes have been described in the general dynamic model of island biogeography (Whittaker et al., 2008), which provides a temporal framework for variations in island features (Lim & Marshall, 2017) such as area, topographical complexity, isolation, and habitat diversity (*a* in EvA; Fig. 5). The model has already been implemented using quantitative methods (Valente, Etienne, & Phillimore, 2014; Borregaard, Matthews, & Whittaker, 2016) and therefore can define the temporal dimension of the abiotic arena in an insular context (Fig. 5).

In the EvA framework, oceanic islands offer a convenient conceptual aspect, in that the abiotic, geographical component of the Evolutionary Arena – the island or archipelago – has discrete boundaries. While many classic studies have analysed insular diversification and disparification (*d* in EvA) in single monophyletic radiations (e.g., Hawaiian silverswords, Baldwin & Sanderson, 1998; Madagascan vangas, Jønsson et al., 2012), a potential of islands is that entire communities resulting from multiple colonisations of the same island can be studied simultaneously, because the abiotic environment is the same for all included taxa with similar dispersal capacity. For instance, by including all terrestrial birds of the Galápagos islands in the same model, Valente, Phillimore, and Etienne (2015) showed that Darwin’s finches have statistically exceptional rates of species diversification (i.e. significantly different from the ‘background’ rates of all terrestrial Galápagos birds). The effect of the phenotype (*c* in EvA; Fig. 5) can be considered on a lineage-specific basis (e.g., Givnish et al., 2009) or, potentially, using a community-level multi-lineage approach, where the effects of given traits are assessed across multiple insular radiations within the same EvA model. Regarding the effect of the biotic interactions on islands (*b* in EvA) it needs to be considered that in most cases, there are precursors to current-day islands that have been eroded to the Pleistocene sea level and are now submerged as guyot seamounts. These previous islands may explain the fact that the evolutionary age of lineages can be older than the respective island where they are endemic today (Pillon & Buerki, 2017). Also, while methods for assessing diversity-dependent effects within single lineages already exist (Rabosky & Glor, 2010; Valente et al., 2015), we currently lack an approach for testing how the interaction of habitat heterogeneity, island size and present diversity can affect all lineages on an island-wide basis.

EvA provides a heuristic framework for the integration of time-dependent model and multi-clade analyses. Once several analyses of clades or archipelagos are available in this framework, it should be possible to combine them to develop a single model for the evolution of diversity within island systems. Furthermore, island disparification and diversification can be compared using the same analytical framework, allowing us to test the hypothesis that they respond to the same factors. The simple EvA framework makes explicit these research question.

#### *Ophrys* biotically driven radiation

It is thought that biotic interactions (*b* in EvA) have been a dominant driver of the radiation of the Mediterranean orchid genus *Ophrys* L., which has produced two parallel adaptive radiations within the last ∼1 Ma. Both of these radiations are characterised by a shift to (mostly) *Andrena* Fabricius solitary bees as highly specific pollinator species, and by rapid disparification of flowers (Paulus & Gack, 1990; Breitkopf, Onstein, Cafasso, Schlüter, & Cozzolino, 2015). In this system, pollinators mediate strong reproductive isolation in the absence of any measurable post-pollination barriers to gene flow among closely related species (Xu et al., 2011; Sedeek et al., 2014). Consequently, pollinators may drive speciation in these two parallel radiations.

The high specificity of the pollinators in the *Ophrys* system is due to the plants’ chemical mimicry of the pollinator females’ sex pheromones (i.e., phenotypic traits, *c* in EvA), which is predominantly mediated by alkenes (Schiestl et al., 1999; Xu, Schlüter, Grossniklaus, & Schiestl, 2012). A simple genetic basis underlies alkene biosynthesis, with only two loci being sufficient to completely change the pollinator-important alkene double-bond profile sensed by insects (Schlüter et al., 2011; Sedeek et al., 2016). Selection on alkene composition, and on loci putatively involved in their biosynthesis (*c* in EvA), is in stark contrast to the rest of the loci in the genome, where abundant polymorphisms are shared across closely related species (Sedeek et al., 2014). Simulations suggest that this simple trait architecture could lead to rapid pollinator-driven divergence (Xu & Schlüter, 2015). Overall, the available data suggest that, given the trait architecture of pollinator attraction, pollinators may be a key factor driving the *Ophrys* radiations (Fig. 6). However, the relative importance of other factors remains unknown. For example, what is the potential contribution to phenotypic variation (i.e. disparification; *d* in EvA) attributable to the highly heterogeneous (dynamic) abiotic environment (*a* in EvA) providing habitats for plants and pollinators (*b* in EvA) during the last million years?

Potentially, proxies for all components are measurable here. Due to the extensive amount of allele sharing, and gene coalescence times frequently predating ‘speciation’ times (establishment of reproductive barriers), diversification (*d* in EvA) among very recent groups of *Ophrys* may best be assessed not by phylogenetic means, but by within-group pairwise estimates among closely related species, either in terms of genetic differentiation (e.g., *F*_ST_) or phenotypic measurements. Among the abiotic measurables (*a* in EvA) would be estimates of habitat fragmentation (also, for example, estimates from biogeographic and niche modelling approaches) over the estimated age of target clades/species groups. Biotic interactions (*b* in EvA) can be represented as matrices of interactions at varying levels; in its simplest form, orchid species vs. pollinator species. Due to the occurrence of (i) parallel use of the same pollinators in different lineages and (ii) repeated use of the same pollinators by allopatric orchid species, such an interaction matrix could initially take the shape of a variance-covariance matrix as in comparative analyses (O’Meara, 2012). Since the importance of different floral traits (including morphology and chemistry) for successful interaction with pollinators can be experimentally measured by quantifying insect responses (e.g., Xu et al., 2012), such an interaction matrix may eventually contain explicit likelihoods of plant/pollinator interactions (cf. pollination probabilities in Xu & Schlüter, 2015). Clade-specific intrinsic effects (*c* in EvA) essentially refer to the complexity of genetic/genomic change needed to effect a change in a relevant trait. This trait change would ideally be based upon mechanistic molecular knowledge that could then be quantified numerically. As a first step in this direction, it may be possible to construct a statistic that summarises trait–gene associations derived from genomic/transcriptomic data on gene polymorphisms and/or expression from phenotyped individuals. Overall, although the data to formally test the components of the EvA framework in the *Ophrys* example are not yet collected, doing so seems at least theoretically straightforward.

EvA adds a comparative structure to the questions developed above (Table 2). It may also provide a semantic to bridge micro- and macroevolution in systems where investigations span different levels, from populations to species and clades.

**Table 2.** Summary of EvA components for the four examples detailed above: worldwide conifers (quantified), Andean *Lupinus*, island radiations and Mediterranean *Ophrys*.

| Case | *d* | *a* | *b* | *c* |
| --- | --- | --- | --- | --- |
| Conifers | Net diversification rate | Potential area size<br>(0.001 % *r*<sup>2</sup>) | Competition<br>(36 % *r*<sup>2</sup>) | Physiological traits<br>(2.2 % *r*<sup>2</sup>) |
| *Lupinus* | Speciation<br>Growth form disparification<br>Genetic differentiation | Extent of páramo<br>Physiographical heterogeneity<br>Flickering connectivity | Low<br>competition (?) | Secondary perennality |
| Islands | Diversification<br>Disparification | Island ontogeny<br>Isolation<br>Fragmentation | Competition for<br>resources | Idiosyncratic (?) |
| *Ophrys* | Speciation | Fragmentation of range | Pollinators | Flower morphology<br>Floral traits, esp. odours<br>Genetics |

## Conclusions

In this review we synthesize the central concepts of the evolutionary diversification literature and present the Evolutionary Arena framework (EvA), a heuristic for exploring the modulators of diversification rates, in terms of the (extrinsic) biotic and abiotic environment and the (intrinsic) traits of the focal lineage (Table 2), i.e. we integrate three comprehensive classes of diversification rate modulators. The framework encourages us to organise knowledge about the factors regulating evolutionary radiations, evolutionary stasis and evolutionary decline of lineages. It does so in a generalized framework and helps recognise missing information. Particularly in the exploratory phases of research, EvA may support the search for more complete explanations of diversification rate variation.

The framework is very flexible, facilitating the incorporation of detailed variables, interactions among the components, changes in the direction of effect of these components, and interpretation of phylogenetic conservatism and trait lability. EvA advocates a multivariate perspective on radiations and can be readily expanded to accommodate increasing levels of complexity, to test for the interactions among variables, or to rank variables according to their relative influence on diversification rates. Whether the specific results are tallied, or whether the predictors are collapsed, will most likely depend on the type of question being asked, and on the power available in the study system. EvA can be formulated as a hypothesis-testing framework to test whether the likelihood of observing the data under a favourite particular model provides better fit than an appropriate null model, or to compare models of varying complexity. The framework may be particularly useful in parameterising data-rich, broad-scale analyses comparing different systems, such as evolutionary radiations of clades across different regions, or between different clades within the same region, for example, the plant radiations in the north Andean páramos. Applying these analyses within the framework allows us to identify the important components that account for differences in diversification rates between clades and regions. The Evolutionary Arena framework thus encourages a more comparative approach to exploring phylogenetic and geographical variation in the correlates and drivers of speciation and extinction.

## Supporting information

Supplemental R script

## Glossary

Adaptation: a trait is an adaptation to a selective regime if it evolved in response to selection by that regime (Gould & Vrba, 1982)
Adaptive zone: a fitness peak in a set of related niches (the adaptive grid or macroevolutionary landscape) that a lineage occupies by virtue of a novel trait(s) that confer fitness in these niches.
Confluence: the sequential coming together of a set of traits (innovations and synnovations), environmental changes, and geographic movements along the branches of a phylogenetic tree (Donoghue & Sanderson, 2015).
Disparification: increase in trait variance in a clade through time. That is, increase in measurable phenotypic differences among taxa, whereas the traits in question may be morphological, anatomical, physiological, genetic, behavioural, etc. Disparification is a characteristic of adaptive radiation.
Diversification: increase in the taxonomic diversity in a clade through time. The diversification rate is defined as speciation *minus* extinction and can thus be negative.
Ecological opportunity: lineage-specific environmental conditions that contain both niche availability and niche discordance, favouring adaptation and promoting diversifying selection within the lineage (adapted from Wellborn & Langerhans, 2015).
Exaptation: a trait that has evolved for one use, and that is later useful for another usage (sometimes deceptively termed ‘pre-adaptation’; adapted from de Vladar et al., 2017). The original definition of Gould and Vrba (1982) is: “features that now enhance fitness but were not built by natural selection for their current role”.
Extrinsic factor: environmental factors such as abiotic and biotic niche parameters, not inherited genetically by the focal lineage.
Intrinsic factor: phenotypic (morphological, physiological) or genetic trait(s), inherited by the focal lineage.
Key event: events that trigger a shift in diversification rates.
Key innovation: new trait which facilitates the occupation of a new adaptive zone, or which breaks an evolutionary constraint; i.e. “phenotypes that allowed a species to interact with the environment in a novel way” (Stroud & Losos, 2016, p 508).
Phenotype: a set of features of an individual that stems from the interactions between genotype and environment.
Radiation: accelerated proliferation of species and/or phenotypes, in the sense of significant increase in the diversification and/or disparification rate compared to background rates (without a shift/ significant rate increase, it is not a radiation but [background] diversification / disparification). Radiation (diversification / disparification) can be combined with an epithet, such as adaptive, geographical, ecological, or genetical (gene flow) to further describe the nature of the evolutionary forces and situations (e.g., ‘sexual radiation’ may refer to radiation driven by sexual selection, or ‘montane radiation’ to radiation in mountains, ‘insular radiation’ to radiations on islands, or simply ‘cichlid radiation’ to refer to a certain lineage, etc.). We refer to biological radiation most generally as ‘**evolutionary radiation**’ as the change in diversification / disparification rates has macro-evolutionary consequences. In addition, there are two prominent concepts that refer to the process underlying the evolutionary radiations:
Adaptive radiation: proliferation of species driven by the evolution of phenotypic (ecological and/or morphological) diversity that can be linked to adaptation to an environment. The environment may act as a modulator, driving (potentially sympatric) speciation and/or slowing extinction.
Geographic radiation: proliferation of species driven by enhanced opportunities for allopatric speciation (reproductive isolation resulting from spatial barriers) in a particular region (modified from Simões et al., 2016). Also referred to as ‘non-adaptive radiation’ (= geographic), or ‘climatic radiation’ when differing climates are thought important. Note that the adaptive and geographic categories are simplified: both adaptive and neutral processes likely play a role in modulating diversification rates in most radiations, but their relative contributions differ. For example, ecological factors may enhance the opportunity for reproductive isolation, and species divergence in adaptive radiations may additionally be promoted by spatial isolation (see main text, Drivers of evolutionary radiations).
Synnovation: interacting combination of traits with a particular consequence (Donoghue & Sanderson, 2015).
Trait: a heritable attribute of evolutionary lineages (genes, individuals, populations, species, clades) that can be observed.
Trigger: Event or situation starting a radiation.

## Data Accessibility Statement

The phylogenetic tree, the data matrix and R scripts used in the analysis of conifers are available at Dryad: doi: 10.5061/dryad.2bvq83bkx (Nürk, Linder, et al., 2019).

## Author Contributions section

HPL and NMN developed research concept, all authors contributed to developing the Evolutionary Arena framework, MJL, NMN and REO analysed the data, CEH, LPF, LV, and PMS contributed the conceptual examples. HPL and NMN wrote the manuscript and all authors contributed critically to the drafts and gave final approval for publication.

## Acknowledgements

We especially thank Seraina Klopfstein, Susanne Renner and Bob Ricklefs for discussions and important comments, Andreas Franzke for commenting on an earlier version of this manuscript, and Isabell Niclas, Stefan Pelka and Bernhard Maier for their help during an ambulatory workshop meeting in August 2018 where the ideas presented here were discussed and developed, and which was supported by a WinUBT (Bayreuth) conference grant to NMN. REO acknowledges the support of the German Centre for Integrative Biodiversity Research (iDiv) Halle-Jena-Leipzig funded by the Deutsche Forschungsgemeinschaft (DFG, German Research Foundation)—FZT 118. SGAF acknowledges the ERC Advanced Grant 741413 Humans on Planet Earth (HOPE).

