## Supplemental R script for "Diversification in evolutionary arenas – assessment and synthesis"

### R script used for 'the Conifer EvA'

*03 Dec 2019*

This script runs through the Conifer case study in Nürk et al. "Diversification in evolutionary arenas — assessment and synthesis" applying the Evolutionary Arena (EvA) framework using phylogenetic generalized least squares (pgls). It relies on the files "dd_Conifers_standardized_EvA_pgls.txt" and "tree455_v2_reduced_for_EvA.tre" to be in the working directory of your R session.

For questions:, or

The script uses the following R-libraries:

```
library(ape)
library(geiger)
library(phytools)
library(phylolm)
library(rr2)
```

The code below loads and adjusts the data, runs a pgls analysis, and calculates the R^2 and the partial r^2 values for the independent variables. Finally, the code for the original plots used in Fig. 3 in the paper is given.

#### Load the Conifers data and the time tree

```
dat <- read.table(file = "dd_Conifers_standardized_EvA_pgls.txt")
phy <- read.nexus("tree455_v2_reduced_for_EvA.tre")
```

#### Adjust data sets

Because monotypic clades don't have a niche evolution rate (calculated between all species within a clade!), NAs are in 'dat' for 25 (monotypic) tips, which we are removing here from 'dat' and 'phy':

```
## remove NAs from dat
dat.r <- dat[!is.na(dat$log_nichEvolRate), ]

## prune phy accordingly
phy.r <- drop.tip(phy, name.check(phy, dat.r)$tree_not_data)

## check for congruency of names/tips
name.check(phy.r, dat.r)
```

```
## [1] "OK"
```

#### Fit PGLS (full model; using phylolm)

```
fit.plm <- phylolm(sqrt_divrate ~ log_nicheArea + expectedComp + log_nichEvolRate,
                   data = dat.r,  phy.r, model = "lambda", lower.bound = 0)
```

##### Summary statistics of full model (fit.plm)

```
( res.plm <- summary(fit.plm) )
```

```
##
## Call:
## phylolm(formula = sqrt_divrate ~ log_nicheArea + expectedComp +
##     log_nichEvolRate, data = dat.r, phy = phy.r, model = "lambda",
##     lower.bound = 0)
##
##    AIC logLik
## -28.93  20.46
##
## Raw residuals:
##       Min        1Q    Median        3Q       Max
## -0.285086 -0.118928 -0.007962  0.074373  0.293818
##
## Mean tip height: 337.0423
## Parameter estimate(s) using ML:
## lambda : 1.247422e-08
## sigma2: 6.401882e-05
##
## Coefficients:
##                    Estimate     StdErr t.value   p.value
## (Intercept)       0.6005521  0.1596667  3.7613 0.0005848 ***
## log_nicheArea    -0.0026438  0.1482564 -0.0178 0.9858681
## expectedComp     -0.5813767  0.1274132 -4.5629 5.394e-05 ***
## log_nichEvolRate  0.1662597  0.1809149  0.9190 0.3640513
## ---
## Signif. codes:  0 '***' 0.001 '**' 0.01 '*' 0.05 '.' 0.1 ' ' 1
##
## Note: p-values are conditional on lambda=1.247422e-08.
```

##### Plot residuals of full model (fit.plm)

```
par(mfrow=c(2,2))
plot(density(fit.plm$residuals))
qqnorm(fit.plm$residuals); qqline(fit.plm$residuals)
plot(fit.plm$fitted.values, fit.plm$residuals)
plot(fit.plm)
```

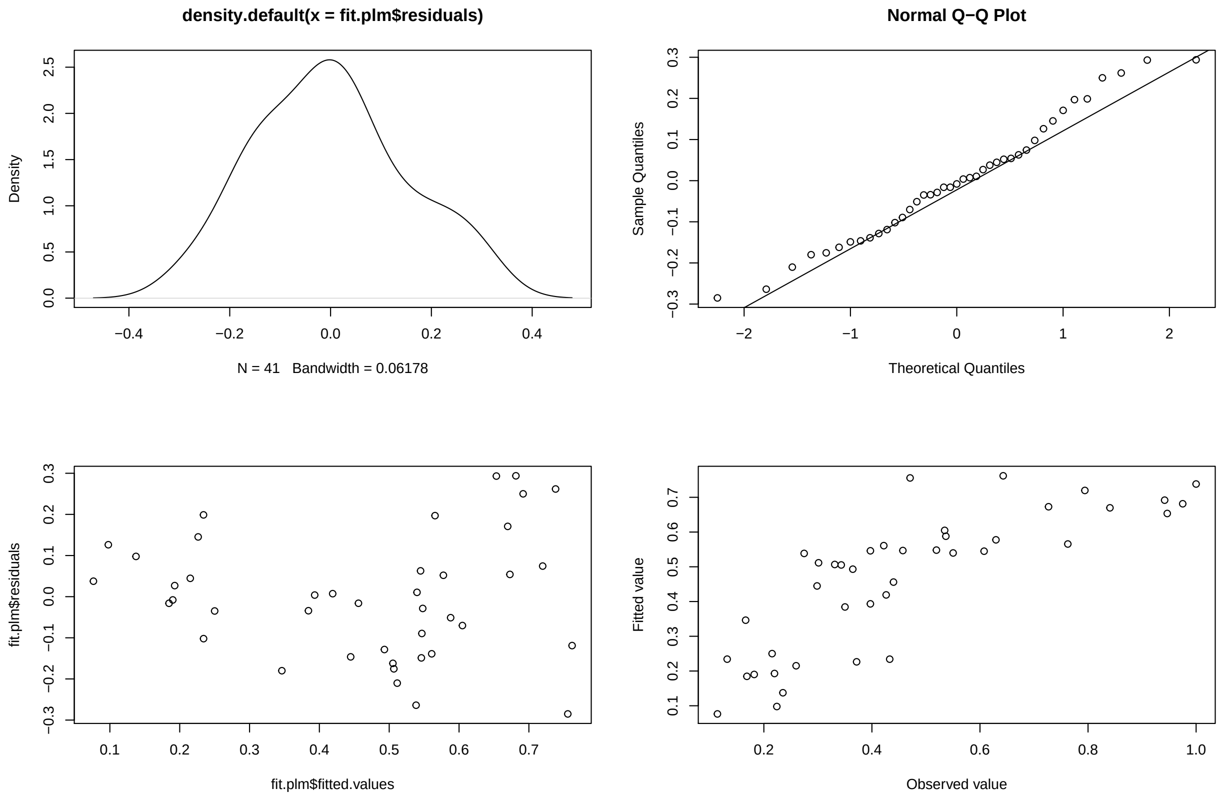


#### Fit partial (reduced) PGLS models (for partial r^2)

```
## 1. Measure the variance explained by predictor (independent) variables
## log_nicheArea (nA)
fit.plm.nA <- phylolm(sqrt_divrate ~ expectedComp + log_nichEvolRate,
                      data = dat.r,  phy.r, model = "lambda", lower.bound = 0)
## expectedComp (eC)
fit.plm.eC <- phylolm(sqrt_divrate ~ log_nicheArea + log_nichEvolRate,
                         data = dat.r,  phy.r, model = "lambda", lower.bound = 0)
## log_nichEvolRate (nE)
fit.plm.nE <- phylolm(sqrt_divrate ~ log_nicheArea + expectedComp,
                         data = dat.r,  phy.r, model = "lambda", lower.bound = 0)

##2. Measure the variance explained by the covariances (variance components)
## phylogeny: "how much var. is explained by the phylo?" (contri. of corrlation structure)
phy.r.x <- compute.brlen(phy.r, method = "Grafen",
                            power = .0001) #construct a star phylogeny ("white noise")
fit.plm.x <- phylolm(sqrt_divrate ~ log_nicheArea + expectedComp + log_nichEvolRate,
                        data = dat.r,  phy.r.x, model = "lambda", lower.bound = 0)
```

##### Calculate the R^2 (for the full model) and partial r^2 (for the independent variables)

```
## full model (fit.plm)
R2(fit.plm, phy = phy.r, resid = FALSE)
##    R2_lik   R2_pred
## 0.6386134 0.6386134
##
## log_nicheArea
( r2.nA <- R2(fit.plm, fit.plm.nA, phy = phy.r, resid = FALSE) )
##       R2_lik      R2_pred
## 8.593904e-06 8.594745e-06
##
## expectedComp
( r2.eC <- R2(fit.plm, fit.plm.eC, phy = phy.r, resid = FALSE) )
##   R2_lik  R2_pred
## 0.360086 0.360086
##
## log_nichEvolRate
( r2.nE <- R2(fit.plm, fit.plm.nE, phy = phy.r, resid = FALSE) )
##     R2_lik    R2_pred
## 0.02231629 0.02231629
##
## phylogeny
( r2.x <- R2(fit.plm, fit.plm.x, phy = phy.r, resid = FALSE) )
##        R2_lik       R2_pred
## -4.531158e-10  1.110223e-16
```

##### Summarize main results

```
## extract estimates (slope, slope_SE, t.val, p.pal)
res.partl.r2 <- as.data.frame(round(res.plm$coefficients, 5))

## add the partial R^2s(pred)
res.partl.r2$percent.partl.r2pred <- round(c(NA, r2.nA[2], r2.eC[2], r2.nE[2]), 5)*100
res.partl.r2
```

```
##                   Estimate  StdErr  t.value p.value percent.partl.r2pred
## (Intercept)        0.60055 0.15967  3.76129 0.00058                   NA
## log_nicheArea     -0.00264 0.14826 -0.01783 0.98587                0.001
## expectedComp      -0.58138 0.12741 -4.56292 0.00005               36.009
## log_nichEvolRate   0.16626 0.18091  0.91899 0.36405                2.232
```

##### Figures

```
## get scale propotional to the slope (~indication of 'mean' effects) of predictor variables
( arrow.size <- round(sqrt(abs(res.partl.r2$Estimate[2:4]*400)), 1)/2 )
```

```
## [1] 0.5 7.6 4.1
```

```r
## plot circular trees with density of the predictor variables nA, eC, nE
## prepare trait vectors
trait.nA <- dat.r$log_nicheArea
names(trait.nA) <- phy.r$tip.label
trait.eC <- dat.r$expectedComp
names(trait.eC) <- phy.r$tip.label
trait.nE <- dat.r$log_nichEvolRate
names(trait.nE) <- phy.r$tip.label
# and for the net diversification rates (this is for illustration purpose mainly,
# but see Sakamoto & Venditti 2018 Biol. Lett. http://dx.doi.org/10.1098/rsbl.2018.0502)
trait.div <- dat.r$sqrt_divrate
names(trait.div) <- phy.r$tip.label

## use phytools' fastAnc in contMap to 'color' branches
phy.nA <- setMap(contMap(phy.r, trait.nA, plot = FALSE), invert = TRUE)
phy.eC <- setMap(contMap(phy.r, trait.eC, plot = FALSE), invert = TRUE)
phy.nE <- setMap(contMap(phy.r, trait.nE, plot = FALSE), invert = TRUE)
phy.div <- setMap(contMap(phy.r, trait.div, plot = FALSE), invert = TRUE)

## plot trees (to get a scale)
op <- par(mfrow=c(2, 2), oma = c(1, 1, 1, 1) + 0.1)
plot.contMap(phy.nE, type = "fan", ftype = c("off", 0.1), type = "fan",
             sig = 1, outline = FALSE, lwd = 2.69)
plot.contMap(phy.div, type = "fan", ftype = c("off", 0.1), type = "fan",
             sig = 1, outline = FALSE, lwd = 2.69)
plot.contMap(phy.nA, type = "fan", ftype = c("off", 0.1), type = "fan",
             sig = 1, outline = FALSE, lwd = 2.69)
plot.contMap(phy.eC, type = "fan", ftype = c("off", 0.1), type = "fan",
             sig = 1, outline = FALSE, lwd = 2.69)
```

```r
## plot circular trees with bars (used in Fig. 4)
op <- par(mfrow=c(2, 2), oma = c(1, 1, 1, 1) + 0.1, mar = c(3, 1.5, 1.5, 3))
plotTree.wBars(phy.nE$tree, trait.nE, type = "fan", lwd = 2.69,
               lend = 0, tip.labels = FALSE, fsize = 0.7,
               method = "plotSimmap", colors = phy.nE$cols,
               col = "grey80", border = 0, width = 30, scale = 40)
text(5, -40, labels =  "c: niche evolution", cex = 2, col = "grey60")
plotTree.wBars(phy.div$tree, trait.div, type = "fan", lwd = 2.69,
               lend = 0, tip.labels = FALSE, fsize = 0.7,
               method = "plotSimmap", colors = phy.div$cols,
               col = "grey80", border = 0, width = 30, scale = 40)
text(5, -40, labels =  "d: diversification", cex = 2, col = "grey60")
plotTree.wBars(phy.nA$tree, trait.nA, lwd = 2,
               lend = 0, tip.labels = FALSE, fsize = 0.7,
               method = "plotSimmap", type = "fan", colors = phy.nA$cols,
               col = "grey80", border = 0, width = 30, scale = 40)
text(5, -40, labels =  "a: abiotic", cex = 2, col = "grey60")
plotTree.wBars(phy.eC$tree, trait.eC, type = "fan", lwd = 2.69,
               lend = 0, tip.labels = FALSE, fsize = 0.7,
               method = "plotSimmap", colors = phy.eC$cols,
               col = "grey80", border = 0, width = 30, scale = 40)
```

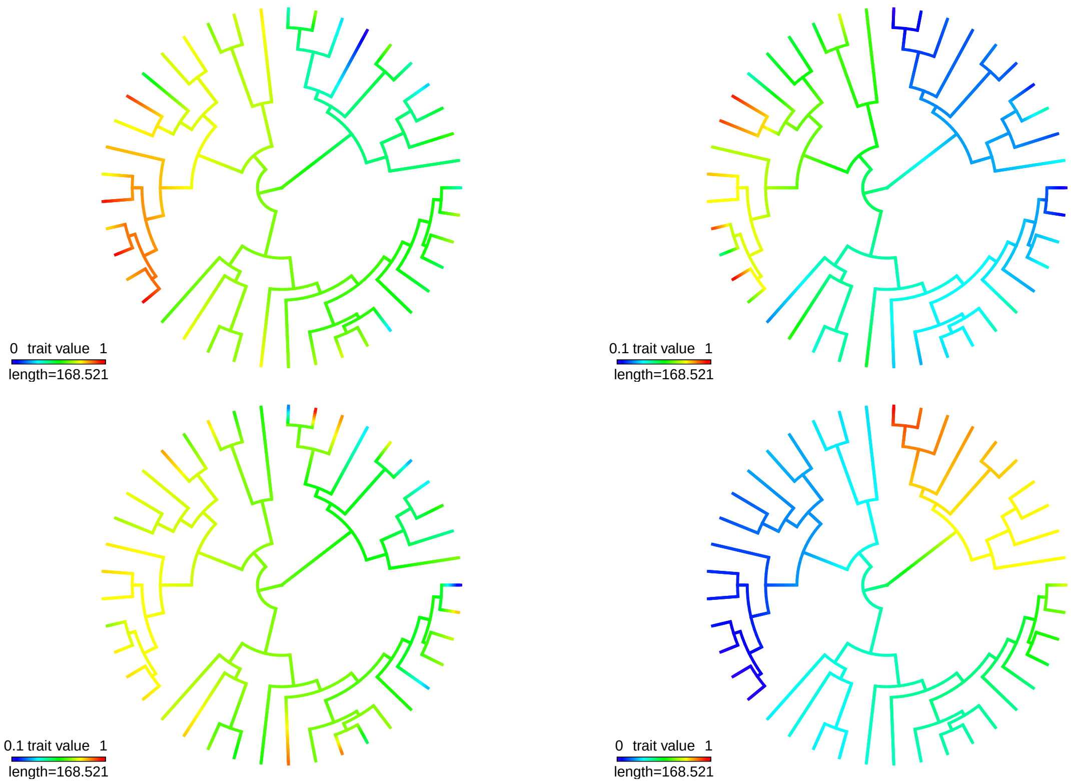


Figure 1: The evolutionary arena of conifers: factor-trees

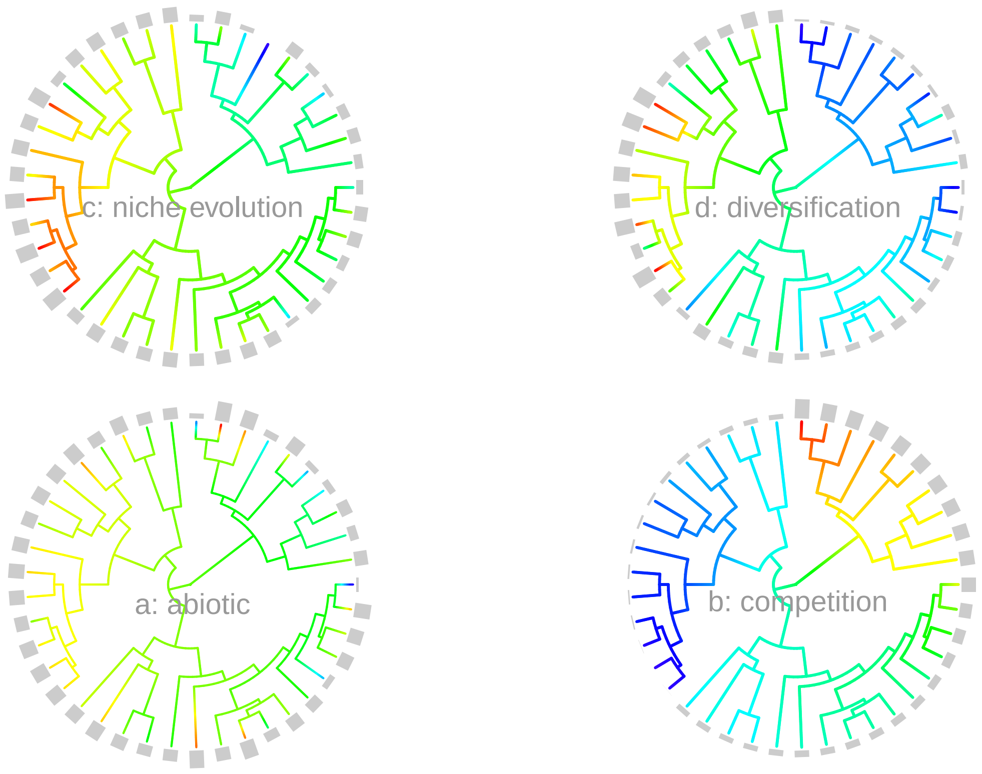


Figure 2: The evolutionary arena of conifers: factor-trees with bars

```
text(5, -40, labels =  "b: competition", cex = 2, col = "grey60")
```

---

---

### ADDITIONAL

#### (not included in the main study):

##### PGLS considering interactions among the predictive components

In the following, the code is given for a full conifer EvA model that also takes interactions among predictor variables (the a, b, and c components) into account

```
fit.plm.ext <- phylolm(sqrt_divrate ~ log_nicheArea * expectedComp * log_nichEvolRate,
                       data = dat.r,  phy.r, model = "lambda", lower.bound = 0)
```

**Summary statistics of full ‘interaction’ model (res.plm.ext)**

```r
( res.plm.ext <- summary(fit.plm.ext) )
```

```
##
## Call:
## phylolm(formula = sqrt_divrate ~ log_nicheArea * expectedComp *
##     log_nichEvolRate, data = dat.r, phy = phy.r, model = "lambda",
##     lower.bound = 0)
##
##    AIC logLik
## -29.13  24.56
##
## Raw residuals:
##      Min       1Q   Median       3Q      Max
## -0.31850 -0.09592  0.03309  0.08546  0.27239
##
## Mean tip height: 337.0423
## Parameter estimate(s) using ML:
## lambda : 0.2885459
## sigma2: 5.741086e-05
##
## Coefficients:
##                                            Estimate  StdErr t.value
## (Intercept)                                 -1.6316  1.0536 -1.5486
## log_nicheArea                                2.6476  1.4285  1.8534
## expectedComp                                 2.0125  1.2747  1.5787
## log_nichEvolRate                             3.6393  1.6862  2.1583
## log_nicheArea:expectedComp                  -2.6816  1.7423 -1.5392
## log_nicheArea:log_nichEvolRate              -4.1412  2.2305 -1.8566
## expectedComp:log_nichEvolRate               -4.2956  2.1804 -1.9701
## log_nicheArea:expectedComp:log_nichEvolRate  4.3751  2.8794  1.5195
##                                            p.value
## (Intercept)                                0.13101
## log_nicheArea                              0.07279 .
## expectedComp                               0.12394
## log_nichEvolRate                           0.03827 *
## log_nicheArea:expectedComp                 0.13330
## log_nicheArea:log_nichEvolRate             0.07231 .
## expectedComp:log_nichEvolRate              0.05727 .
## log_nicheArea:expectedComp:log_nichEvolRate 0.13817
## ---
## Signif. codes:  0 '***' 0.001 '**' 0.01 '*' 0.05 '.' 0.1 ' ' 1
##
## Note: p-values are conditional on lambda=0.2885459.
```

```r
## R^2 for the full extenbded model (fit.plm.ext)
R2(fit.plm.ext, phy = phy.r, resid = FALSE)
```

```
##    R2_lik   R2_pred
## 0.7041182 0.7155024
```

##### Plot residuals of res.plm.ext

```
par(mfrow=c(2,2))
plot(density(fit.plm.ext$residuals))
qqnorm(fit.plm.ext$residuals); qqline(fit.plm.ext$residuals)
plot(fit.plm.ext$fitted.values, fit.plm.ext$residuals)
plot(fit.plm.ext)
```

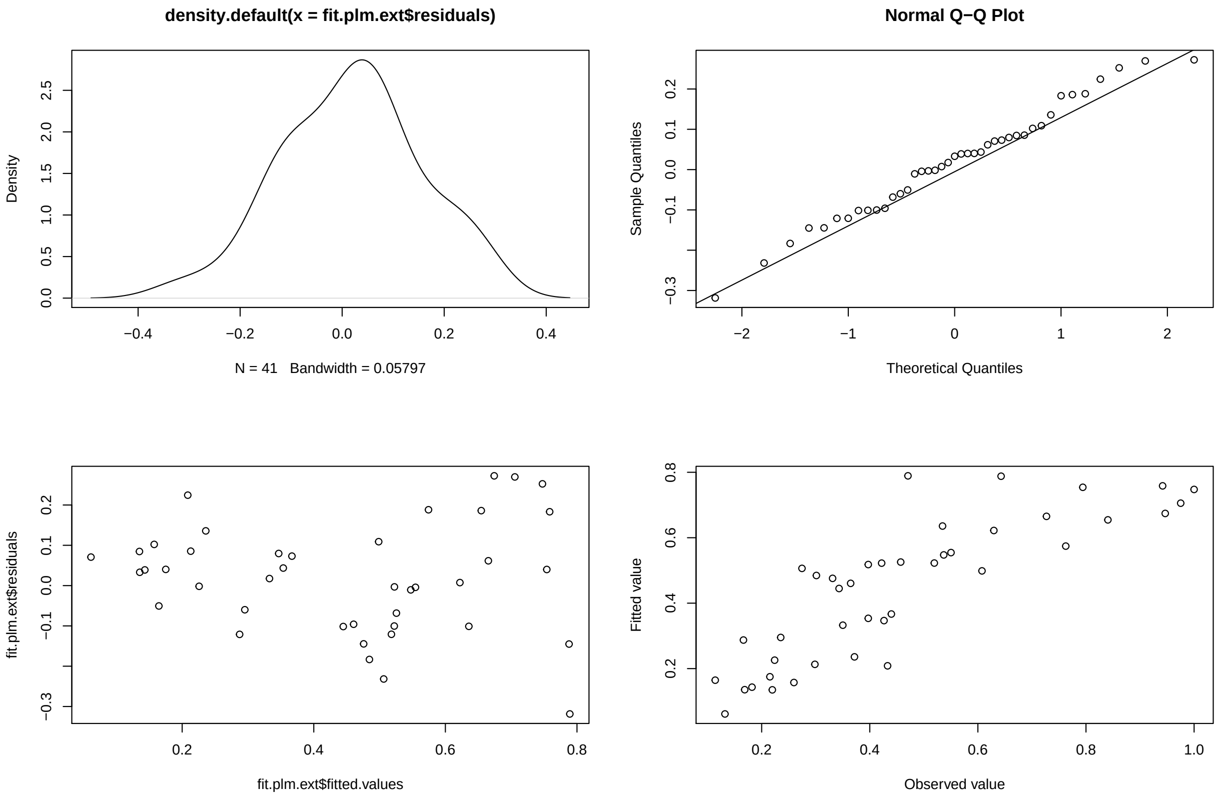


```
plot.phylo(phy.r, type = "fan", cex = .69, no.margin = TRUE)
```

#END

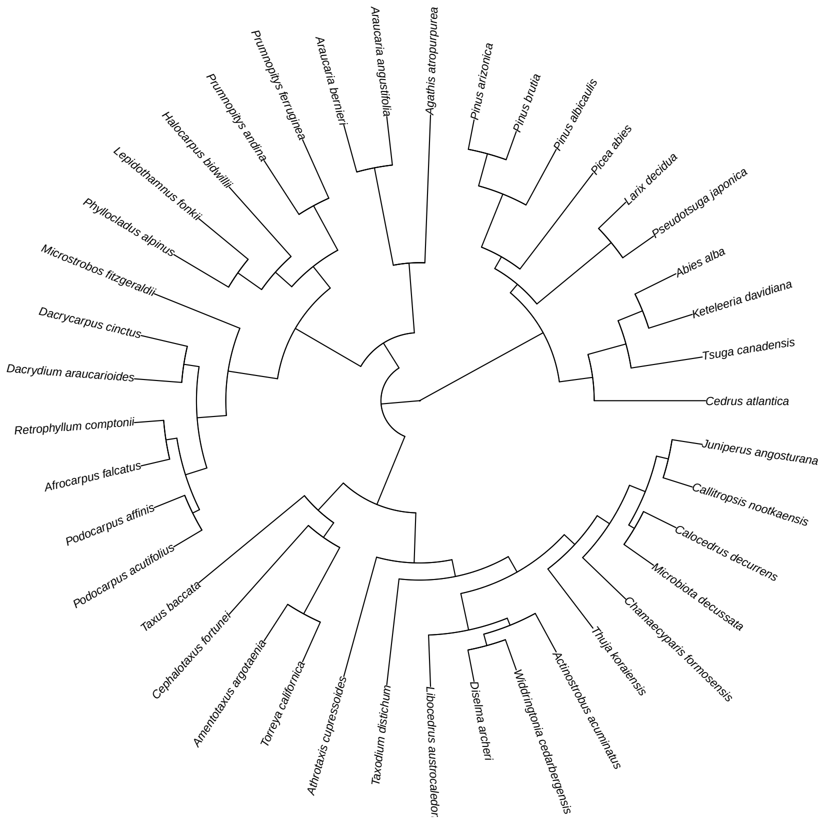


Figure 3: Phylogeny of the conifer clades used in the study, with species names representing the clade
